# A dynein-driven nucleokinesis program enables neural crest migration through confined tissues *in vivo*

**DOI:** 10.64898/2026.06.09.730088

**Authors:** Soraya Villaseca, Minjia Pan, Cheng-Yu Huang, Chengxi Zhu, Marie de la Burgade, Atrin Hamidazdeh, Dina El-Zohiry, Zain Alhashem, Hanna-Maria Häkkinen, Sarah Baxendale, Mar Marzo, Martin O. Lenz, Tanya T. Whitfield, Elena Scarpa

**Affiliations:** Department of Physiology, Development and Neuroscience, University of Cambridge, Downing Street, Cambridge, CB2 3DY, United Kingdom; Cambridge Advanced Imaging Centre, University of Cambridge, Downing Street, Cambridge, CB2 3DY, United Kingdom; School of Biosciences, Bateson Centre for Disease Mechanisms, and Neuroscience Institute, University of Sheffield, Sheffield, S10 2TN, United Kingdom

**Author notes:** these authors contributed equally to the work (Minjia Pan, Cheng-Yu Huang).

## Abstract

Cell migration through confined tissue environments requires precise coordination between the nucleus and the cytoskeleton. Translocation of the nucleus through small pores is a limiting step in confined cell migration. Work in immune and cancer cells has illuminated a key role for actin polymerization and Myosin-II contractility in facilitating nucleus translocation through confinement. However, the microtubule cytoskeleton can also adaptively stabilise under confinement. Microtubule-dependent nucleokinesis constitutes an evolutionarily conserved cellular program ensuring directional nucleus translocation by deployment of cytoplasmic dynein motors. However, whether microtubule-driven nucleokinesis can operate during cell migration through confinement remained so far unexplored.

The zebrafish neural crest (NC) provides a unique *in vivo* system to address this question. NC cells are a highly migratory, multipotent precursors of the vertebrate peripheral nervous system, which traverse diverse environments along the anterior–posterior axis of the embryo. In the head, cranial neural crest (cNC) cells migrate through loosely organised tissues, whereas trunk neural crest (tNC) cells navigate narrowly confined tissue spaces.

Here, we show that tNC cells engage a microtubule motor-driven program of nuclear deformation and translocation during confined migration *in vivo*. Under confinement, tNC cells reorganize their microtubules from a perinuclear meshwork into a polarized, centrosome-associated bundle positioned ahead of the nucleus. Laser ablation reveals that this microtubule array actively pulls and deforms the nucleus. Using pharmacological and genetic approaches, we discover that dynein-dependent pulling forces enable nucleus translocation through confinement. Strikingly, this process occurs independently of Rho/ROCK/myosin II–mediated contractility, revealing a novel neuronal-like mode of nucleokinesis operating in confined tissue environments.

Together, our findings uncover a conserved nuclear translocation program deployed during neural crest migration and suggest that microtubule-based nucleokinesis represents a fundamental strategy for navigating tissue confinement *in vivo* during both central and peripheral nervous system development.

## Introduction

Nucleokinesis, the active repositioning of the nucleus within a cell, is essential for correct cell and tissue morphogenesis. For example, it is important for fungal mitosis and meiosis (Morris et al., 1998, Bone and Starr, 2016), for correct distribution of nuclei within muscles (Cadot et al., 2015), and for central nervous system development (Bone and Starr, 2016).

Nuclear positioning often depends on the microtubule cytoskeleton and associated microtubule motors, such as cytoplasmic dynein and kinesins (Bone and Starr, 2016). Dynein is a minus end directed microtubule motor, and key dynein subunits were first discovered by investigating nuclear translocation in fungi (Xiang et al, 1994). Dynein function in nuclear positioning is strikingly conserved across evolution (Bone and Starr, 2016, Morris et al., 1998). In the context of cell migration, nuclear positioning is tightly regulated when cells navigate physically constrained environments, as cells need to translocate the nucleus, the largest organelle of a cell, through small pores. However, in both amoeboid immune cells (Kroll et al., 2023) and in cancer cells migrating through *in vitro* confinement (Thomas et al., 2015, Keys et al., 2024, Ju et al., 2024), nuclear translocation has been linked primarily to actomyosin-driven forces. Many cell types migrating *in vivo* do not conform to a single migratory mode, but instead display plasticity, transitioning between mesenchymal and amoeboid-like behaviours depending on tissue architecture and confinement (Graziani et al., 2022; Barzegar et al., 2024; Olson et al., 2026). Whether the microtubule/dynein axis contributes to nuclear translocation in confined cell migration remains unexplored.

Here, we address these questions *in vivo* using zebrafish neural crest cells (NCs). These highly migratory cells are the multipotent precursors of the peripheral nervous system. Neural crest cells traverse distinct mechanical landscapes during embryogenesis, ranging from relatively unconfined collective migration in the cranial region to highly restricted inter-tissue spaces in the trunk (Hakkinen et al., 2025). Spatial heterogeneity exposes neural crest cells to varying degrees of confinement while maintaining a common cellular identity (Hakkinen et al., 2025). Moreover, neural crest migration has been shown to depend on microtubules (Moore et al., 2013; Villaseca et al., 2025), raising the possibility that microtubule-based mechanisms may regulate nuclear translocation in these cells. We show that nucleokinesis *in vivo* is not exclusively driven by actomyosin contractility under confinement. Instead, it relies on a microtubule/dynein-based mechanism analogous to that described in central nervous system neurons. By comparing cranial and trunk neural crest cells, we uncover a confinement-dependent mode of nuclear translocation in which microtubules and dynein, which localises at the nuclear envelope in confined trunk neural crest cells, generate pulling forces to deform and reposition the nucleus. These findings reveal a conserved, yet context-dependent, mechanism of nucleokinesis that integrates cytoskeletal dynamics with the physical constraints of the tissue environment.

## Results

### Nucleokinesis occurs during *in vivo* confinement independently of membrane extension and retraction

To investigate how cells navigate physical constraints *in vivo*, we used zebrafish neural crest (NC) cells as a model of physiologically confined cell migration. NC cells are highly migratory and, depending on their anteroposterior location within the embryo, migrate either in non-confined environments (Figure 1A) or under strong physical confinement (Figure 1B) (Hakkinen et al., 2025).

**Figure 1.**
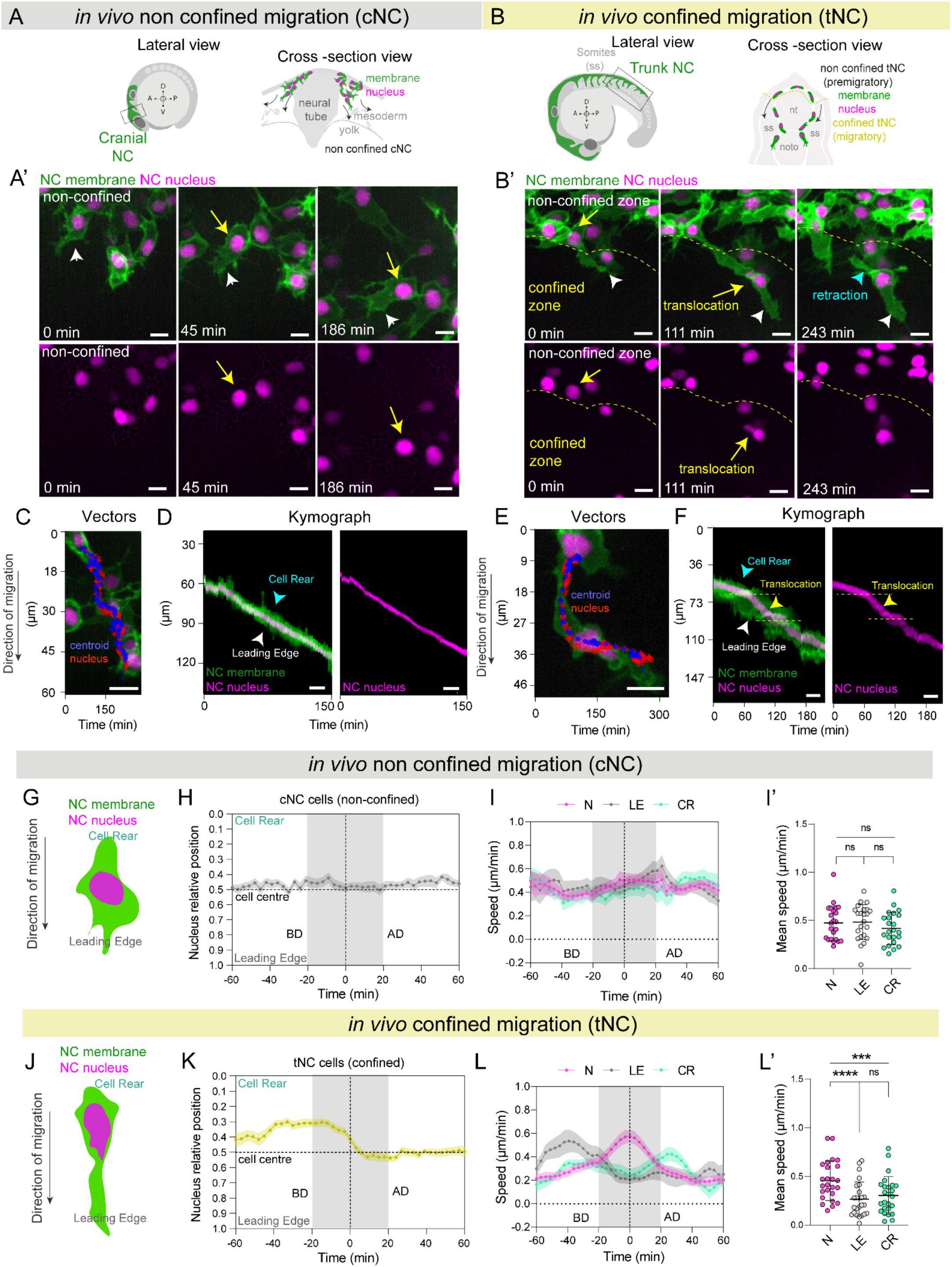
Confined migration uncouples nucleokinesis from cell edge dynamics *in vivo*. (A) Schematic representation of cranial neural crest (cNC) cell migration *in vivo* in a 16-somite stage embryo, shown in lateral and cross-sectional views, illustrating the non-confined environment. cNC cells are depicted in green (membrane) with magenta nuclei, whereas surrounding tissue is showed in grey. (A′) Representative time series of cNC cells migrating from the dorsal to the ventral side of the embryo. White arrowheads indicate membrane dynamics, while yellow arrows indicate nuclear movement. Scale bar, 10 µm. (B) Schematic representation of trunk neural crest (tNC) cell migration *in vivo* in a 20-somite stage embryo, shown in lateral and cross-sectional views, highlighting confinement imposed by surrounding tissues (somites, ss; neural tube, nt; notochord, noto). Cells transition from a non-confined premigratory region to a confined migratory region. (B’) Representative time series of tNC cells migrating from the dorsal to the ventral side of the embryo. White arrowheads indicate membrane dynamics and yellow arrows indicate entry into the confined region and nucleus translocation. Scale bar, 10 µm. (C, E) Schematics summarizing cell migration and the quantitative analysis pipeline for cNC (C) and tNC (E) cells. Cell rear, nuclear centroid, and leading edge positions were tracked over time. Red arrows indicate the direction of nuclear movement and blue dots indicate the cell centroid. Scale bars, 10 µm. (D, F) Representative kymographs of membrane (green) and nuclear (magenta) intensity profiles aligned to the nuclear centroid along the migration axis for cNC (D) and tNC (F) cells. Cell rear dynamics are indicated by cyan arrowheads and leading-edge dynamics by white arrowheads. Nuclear translocation is highlighted in yellow. Scale bars, 10 µm. (G, J) Schematics illustrating the definition of cell geometry during migration in non-confined cNC (G) and confined tNC (J) cells. Cell rear (cyan) was defined opposite to the migration direction, and the leading edge (grey) at the front of the cell. (H, K) Relative nuclear position within the cell over time in cNC (H) and tNC (K) cells, aligned to the point of maximum nuclear deformation (time = 0), defined as the time point of minimum nuclear circularity. Relative nuclear position was calculated as:

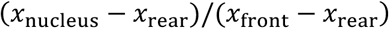 Values approaching 0 indicate rear localization, values near 0.5 indicate central positioning, and values approaching 1 indicate localization toward the leading edge. The grey shaded region indicates the period surrounding nuclear translocation. BD, before nuclear deformation; AD, after nuclear deformation. cNC: *n* = 23 cells from 4 embryos across 3 experiments. tNC: *n* = 25 cells from 6 embryos across 5 experiments. Width of ribbon represents SEM. (I, L) Mean nuclear speed (magenta) compared with leading-edge speed (grey) and cell rear speed (cyan) over time in cNC (I) and tNC (L) cells, aligned to maximum nuclear deformation (time = 0). Data span 60 min before and after alignment. In cNC cells, nuclear and cell edge movements remain coordinated, whereas in tNC cells nuclear movement becomes uncoupled from cell edge dynamics during the nuclear translocation period. BD, before nuclear deformation; AD, after nuclear deformation. Error bars represent SEM. (I’) Statistical comparison of mean nuclear speed, leading-edge speed, and cell rear speed during the nuclear translocation window (−20 to +20 min relative to maximum nuclear deformation) in cNC cells. A one-way repeated measures ANOVA was performed to compare measurements within the same cells. This revealed no significant differences between nuclear and leading-edge speed (N vs LE, *p* = 0.9006), nuclear and cell rear speed (N vs CR, *p* = 0.2240), or leading-edge and cell rear speed (LE vs CR, *p* = 0.1609). *n* = 23 cells from 4 embryos across 3 experiments. (L’) Statistical comparison of mean nuclear speed, leading-edge speed, and cell rear speed during the equivalent analysis window (−20 to +20 min relative to maximum nuclear deformation) in tNC cells. A Friedman test with multiple comparisons was conducted. Nuclear speed differed significantly from both leading-edge speed (N vs LE, *p* < 0.0001) and cell rear speed (N vs CR, p = 0.0004), whereas no significant difference was observed between leading-edge and cell rear speeds (LE vs CR, p = 0.6093). *n* = 25 cells from 6 embryos across 5 experiments.

To characterise nuclear and membrane dynamics during *in vivo* migration, we carried out *in vivo* live imaging using Sox10:mG embryos (Richardson et al., 2016), in which neural crest cell membranes and nuclei are fluorescently labelled (Figure 1A-B′, Supplementary Movie 1). We have previously observed that cranial neural crest (cNC) cells display round shaped nuclei (Hakkinen et al., 2025) (Figure 1A-A’, Supplementary Movie 1, cNC, yellow arrow). By contrast, trunk neural crest (tNC) cells migrate through highly confined spaces (∼3 μm in width), where the nucleus undergoes pronounced shape changes during migration (Hakkinen et al., 2025) (Figure 1B-B’, Supplementary Movie 1, tNC, yellow arrow).

By segmenting membrane and nucleus shapes, we extracted morphometrics for cNC and tNC nuclei and cell shapes. By aligning cell shape metrics to the timepoint of minimum nucleus circularity (maximum nucleus deformation) (Hakkinen et al., 2025), we found that both nucleus (Figure S1A, magenta curve) and cell shapes of non-confined cNCs remained relatively constant throughout migration (Figure S1A-B’, green curve). On the other hand, for confined tNC, we observed a marked decrease in nuclear circularity (Figure S1C, magenta curve), whilst overall cell shape circularity remained unchanged (Figure S1C-C’, green curve). In contrast with cell circularity (Figure S1C’), tNC cell shape elongation is high before nucleus deformation and significantly reduces after nucleus deformation, suggesting cell contraction (Figure S1D-D’, green curve).

To track the movement of the nucleus relative to the cell leading edge and rear boundaries, we analysed membrane dynamics relative to nuclear motion by extracting 1D intensity profiles along the cell’s direction of travel (Figure 1C, 1E). The membrane boundaries were then mapped back to the 1D spline coordinate system, allowing for the construction of kymographs that represent the relative expansion and contraction of the cell body along the direction of cell migration (Figure 1D, 1F, S1E-E’, G-G’). In non-confined cNC cells, nuclear movement remained tightly coupled to membrane dynamics (Figure 1C,D, S1E-F). The nucleus centroid remained close to the cell centroid (Figure 1G,H) and the nucleus moved in coordination with membrane protrusion and retraction (Fig. 1I,I’). Cross-correlation analysis of nuclear speed versus leading edge (Figure S1I, grey dots) or cell rear speed (Figure S1I, cyan dots) shows that nucleus and cell membrane movement are closely coupled in non-confined cNCs (Supplementary Movie 1, cNC, arrowheads).

By contrast, nuclear movement was uncoupled from both the leading edge and the cell rear in confined tNC cells (Figure 1F, arrowheads). Measurement of the nucleus centroid position relative to the cell centroid (Figure 1J) highlights an *intracellular* translocation of the nucleus (Figure 1K, Supplementary Movie 1, tNC, yellow arrow).

Indeed, comparison of nuclear speed against the speed of cell rear and cell front (Figure 1L, L’) shows that the nuclear speed significantly increases at the time of maximum nucleus deformation in tNCs. Leading edge speed decreases concomitantly with the nuclear translocation phase (Figure 1L, gray curve), whilst cell rear movement occurs after nuclear translocation (Figure 1L, cyan curve, Supplementary Movie 1, tNC, cyan arrowhead). In addition, nuclear translocation speed is significantly faster than cell rear speed and leading edge speed (Figure 1L-L’, magenta curve). Cross-correlation analysis of nuclear speed versus leading edge (Figure S1I’, grey dots) or cell rear speed (Figure S1I’, cyan dots) shows that nucleus speed is uncoupled from cell membrane speed.

Together, this shows that confined tNCs, but not non-confined cNCs, undergo intracellular nuclear translocation during their *in vivo* developmental migration.

### Actomyosin contractility is not required for tNC nuclear translocation through confinement

Previous *in vitro* studies using confined cancer cells have proposed that intracellular nuclear translocation can result from a combination of pulling forces at the front and pushing forces generated by actomyosin contraction at the rear (Stöberl et al., 2024). In these systems, actomyosin contractility at the rear cortex is required to drive nuclear translocation through constrictions (Thomas et al., 2015, Mistriotis et al., 2019, Keys et al., 2024). However, in our *in vivo* model, we observed a markedly different behaviour. During confined migration, trunk neural crest (tNC) cells do not exhibit rear contraction prior to nuclear translocation (Figure 1B’,F, L). Instead, the leading edge extends first, followed by a fast translocation of the nucleus and by delayed retraction of the cell rear, indicating an asynchronous process (Figure 1B’,F, L, Supplementary Movie 1).

To investigate Myosin II localisation, we generated a neural-crest-specific sox10:Myosin-light-chain-emGFP transgenic zebrafish and carried out live imaging of tNCs initiating confined migration (Figure 2A, Supplementary Movie 2). *In vivo* confined tNC cells displayed weak Myosin localization at the cell leading edge and at the cell-cell contacts (Figure 2B, pseudo colour panels, arrows, Supplementary Movie 2: arrowheads) and a stronger accumulation of Myosin at the cell rear following nuclear translocation (Figure 2B, pseudo colour panels, arrowheads, Supplementary Movie 2: arrowhead). To ask whether Myosin-II contractility is required for nucleus translocation of tNCs, we inhibited ROCK using Y-27632 (Figure 2C,D, Supplementary Movie 3). Analysis of nuclear morphometrics revealed that ROCK inhibition did not impair nuclear shape changes (Figure 2E, F). Analysis of leading edge and cell rear dynamics relative to the nucleus (Figure 2G, H) reveals that neither intracellular nucleus translocation (Figure 2I, Supplementary Movie 3: yellow arrow) nor nucleus speed are affected by ROCK inhibition (Figure 2J-L, S2D). Notably, the nucleus remains at the cell centre, rather than returning to the cell rear, after its translocation (Figure 2I black arrow, Figure S2A).

**Figure 2.**
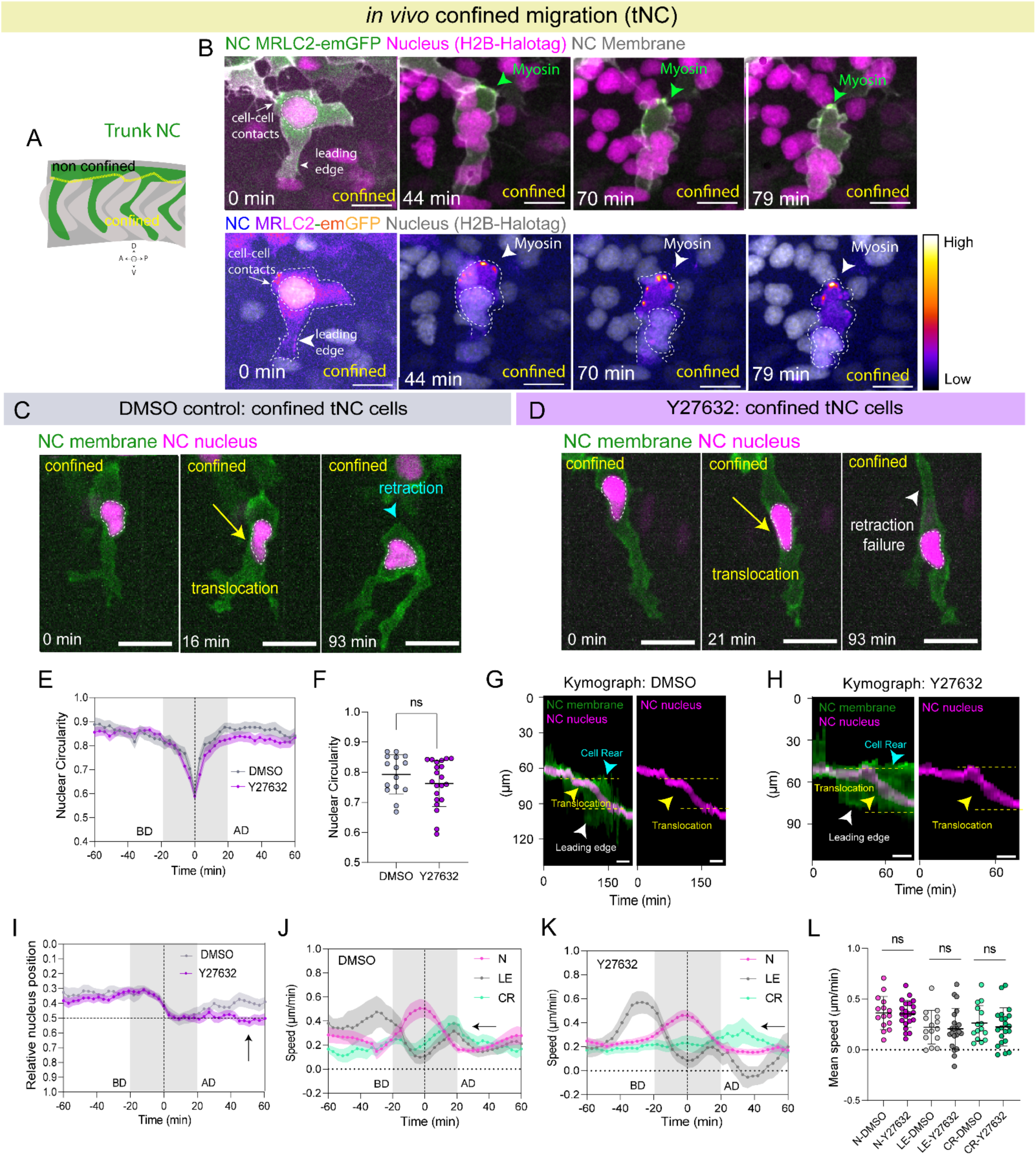
Nucleokinesis and nuclear deformation occur independently of ROCK-mediated actomyosin contractility *in vivo*. (A) Schematic of the mid-trunk neural crest region (between the 7th and 10th somites), highlighting the transition from a non-confined (premigratory) to a confined (migratory) environment. (B) Representative time series of a single tNC cell transitioning from a non-confined to a confined region. Images were acquired from Sox10:MRLC2-emGFP / Sox10:mRFP transgenic zebrafish embryos (green and grey, respectively) injected with H2B-halo mRNA to label nuclei (magenta). Pseudocolour intensity reveals mild enrichment of MRLC2-emGFP at cell–cell contacts and the leading edge, and pronounced accumulation at the cell rear coinciding with membrane retraction (arrowheads). Scale bar, 10 µm. (C, D) Representative time series of tNC cells undergoing confinement in embryos treated with DMSO (C) or 50 µM Y27632 (D) to inhibit ROCK-dependent contractility. Yellow arrows indicate nuclear translocation, cyan arrowheads indicate membrane retraction, and white arrowheads indicate retraction failure following Y27632 treatment. Nuclear deformation and translocation persist despite impaired membrane retraction. Scale bar, 10 µm. (E) Mean nuclear circularity over time in DMSO- and Y27632-treated embryos, aligned to the point of maximum nuclear deformation (time = 0), defined as the minimum nuclear circularity point. Data span 60 min before and after alignment. BD: Before deformation, AD: After deformation. DMSO: *n* = 15 cells from 4 embryos across 3 experiments. Y27632: *n* = 21 cells from 8 embryos across 4 experiments. Error bars represent SEM. (F) Quantification of mean nuclear circularity during the nuclear translocation window (−20 to +20 min relative to maximum nuclear deformation) in DMSO- and Y27632-treated embryos. Each dot represents one cell. No significant difference was observed between conditions (two-tailed Mann–Whitney test, *p* = 0.5606). DMSO: *n* = 15 cells from 4 embryos across 3 experiments. Y27632: *n* = 21 cells from 8 embryos across 4 experiments. (G, H) Representative kymographs of membrane and nuclear intensity profiles aligned to the nuclear centroid along the migration axis in DMSO (G) and Y27632 (H) conditions. Cell rear dynamics are indicated by cyan arrowheads and leading-edge dynamics by white arrowheads. Nuclear translocation events are indicated by yellow arrowheads. Scale bar, 10 µm. (I) Relative nuclear position within the cell over time in DMSO-treated (grey) and Y27632-treated (purple) embryos, aligned to maximum nuclear deformation (time = 0). Values approaching 0 indicate rear localization, values near 0.5 indicate central positioning, and values approaching 1 indicate localization toward the leading edge. The grey shaded region indicates the nuclear translocation period. Black arrows indicate maintenance of a central nuclear position following ROCK inhibition. BD, before deformation; AD, after deformation. DMSO: *n* = 15 cells from 4 embryos across 3 experiments. Y27632: *n* = 21 cells from 8 embryos across 4 experiments. Error bars represent SEM. (J, K) Mean nuclear speed (N, magenta), leading-edge speed (LE, grey), and cell rear speed (CR, cyan) over time in DMSO (J) and Y27632-treated (K) embryos, aligned to maximum nuclear deformation (time = 0). Data span 60 min before and after alignment. The grey shaded region indicates the analysis window (−20 to +20 min). In both conditions, nuclear movement remains uncoupled from cell edge dynamics during nuclear translocation. Black arrows indicate re-coordination of leading-edge and cell rear dynamics after nuclear translocation in DMSO-treated cells, whereas Y27632 treatment disrupts coordination between cell boundaries. DMSO: *n* = 15 cells from 4 embryos across 3 experiments. Y27632: *n* = 21 cells from 8 embryos across 4 experiments. Error bars represent SEM. (L) Statistical comparison of mean nuclear, leading-edge, and cell rear speeds during the analysis window (−20 to +20 min relative to maximum nuclear deformation) in DMSO- and Y27632-treated embryos. One-way ANOVA with multiple comparisons revealed no differences between DMSO and Y27632 conditions for nuclear speed (*p* > 0.9999), leading-edge speed (*p* = 0.9997), or cell rear speed (*p* = 0.9894). DMSO: *n* = 15 cells from 4 embryos across 3 experiments; Y27632: *n* = 21 cells from 8 embryos across 4 experiments.

Quantitative analysis of cell membrane dynamics relative to nuclear motion revealed that, in contrast with control cells, Y-27632 treated cells did not reduce their elongation after nucleus translocation, suggesting an impaired cell rear retraction (Figure S2B-C’, Supplementary Movie 3: arrowhead). Nucleus speed remains higher than leading edge and cell rear speed at the maximum deformation point (Figure 2J-L) and cross-correlation analysis showed that the nucleus speed remains uncoupled from the cell front and cell rear upon inhibition of contractility (Figure S2G). Leading edge speed is not significantly reduced after nuclear translocation (Figure S2E, black arrow), and, whilst cell rear movement following nucleus translocation appears delayed (Supplementary Movie 3: arrowhead), the speed of the cell rear was not significantly affected (Figure S2F). Together, these results suggest that, whilst ROCK activity might help coordinate leading edge extension and cell rear retraction in tNC, nuclear translocation during confined migration *in vivo* occurs independently of actomyosin-driven membrane dynamics. Thus, another intracellular force-generating mechanism likely drives nucleokinesis of tNCs under physiological confinement.

### Microtubules reorganize into a polarized bundle positioned ahead of the nucleus during *in vivo* confined tNC migration

During neuronal migrations of the mammalian neocortex, nucleokinesis is driven by microtubule-dependent forces, where the nucleus is pulled towards the centrosome through dynein activity (Solecki et al., 2006, Tsai et al., 2007, Umeshima et al., 2018, Wang et al., 2009).

In other systems, such as myeloid progenitors, microtubules can generate strong nuclear deformations (Biedzinski et al., 2019). Similarly, during *Drosophila* embryogenesis, microtubules deform nuclei during cellularisation (Schulze et al., 2009). Since we observe striking nuclear deformations associated with nuclear translocation (Figure 1B’, S1C, Supplementary Movie 1), we hypothesised that microtubules may actively contribute to nuclear deformation in this context. Microtubules are highly dynamic polymers that reorganise in response to mechanical constraints (Li et al., 2023). *In vitro*, confined cells assemble a perinuclear network of stable microtubules (Li et al., 2023; Hunter et al., 2025, Ju et al., 2024). However, whether a similar microtubule reorganisation occurs *in vivo* during physiological cell migration remains unknown.

To address this, we analysed microtubule organisation in NC cells *in vivo* using StableMark (Figure 3A, Supplementary Movie 4), a live reporter of stable microtubules (Jansen et al., 2023) or EB3-GFP (Figure 3F, Supplementary Movies 5 and 6), a plus-end tracking protein that labels growing microtubule ends (Stepanova et al., 2005, Moore et al., 2015, Villaseca et al., 2025), by driving their expression under the tissue-specific sox10 promoter (Dutton et al., 2008, Alhashem et al., 2021). In non-confined cNC cells, live imaging of Stable Mark revealed that microtubules organise in a perinuclear mesh-like structure formed by stable microtubules surrounding the nucleus (Figure 3B,C, Supplementary Movie 4, cNC: arrowheads). In contrast, *in vivo* confined tNC cells displayed a striking reorganisation of microtubules, which redistributed from a perinuclear network into a prominent bundle (Figure 3D, 3H, Supplementary Movie 4, tNC: arrowhead, Supplementary Movie 5 tNC: arrowhead). The bundle is constituted of stable microtubules (Figure 3D), and is positioned at the front of the nucleus along the direction of migration (Figure 3D,E). These observations suggest that a subset of microtubules might be stabilised under physical confinement (Li et al., 2023, Ju et al., 2024). To test this quantitatively, we carried out high-time-resolution tracking of EB3-GFP microtubules plus-ends. Comparison between non-confined and confined neural crest cells indeed revealed significant differences in microtubule dynamics. Non-confined cNC displayed non-polarised comet orientation (Figure 3G, 3I, Figure S3A, Supplementary Movie 5), whilst confined tNCs cells exhibited highly oriented comet growth towards the cell leading edge (Figure 3H,3J, Figure S3A, Supplementary Movie 5, tNC: arrowhead). Confined tNCs also show a marked increase in microtubule growth length and growth speed, in comparison with non-confined cNC cells (Figure 3K,L). EB3-GFP comets also exhibited a slightly increased lifetime in confined tNC cells (Figure 3M). Subcellular comparison of perinuclear versus leading edge EB3 dynamics (Figure S3B, C) revealed that both growth length and growth speed increase for perinuclear and leading edge microtubules under confinement (Figure S3D-D’, E-E’). By contrast, comet lifetime was significantly increased at the leading edge (Figure S3F’) but not at the perinuclear area of confined tNC (Figure S3F) in comparison with non-confined cNC, revealing longer lived microtubules specifically at the leading edge of confined NC cells.

**Figure 3.**
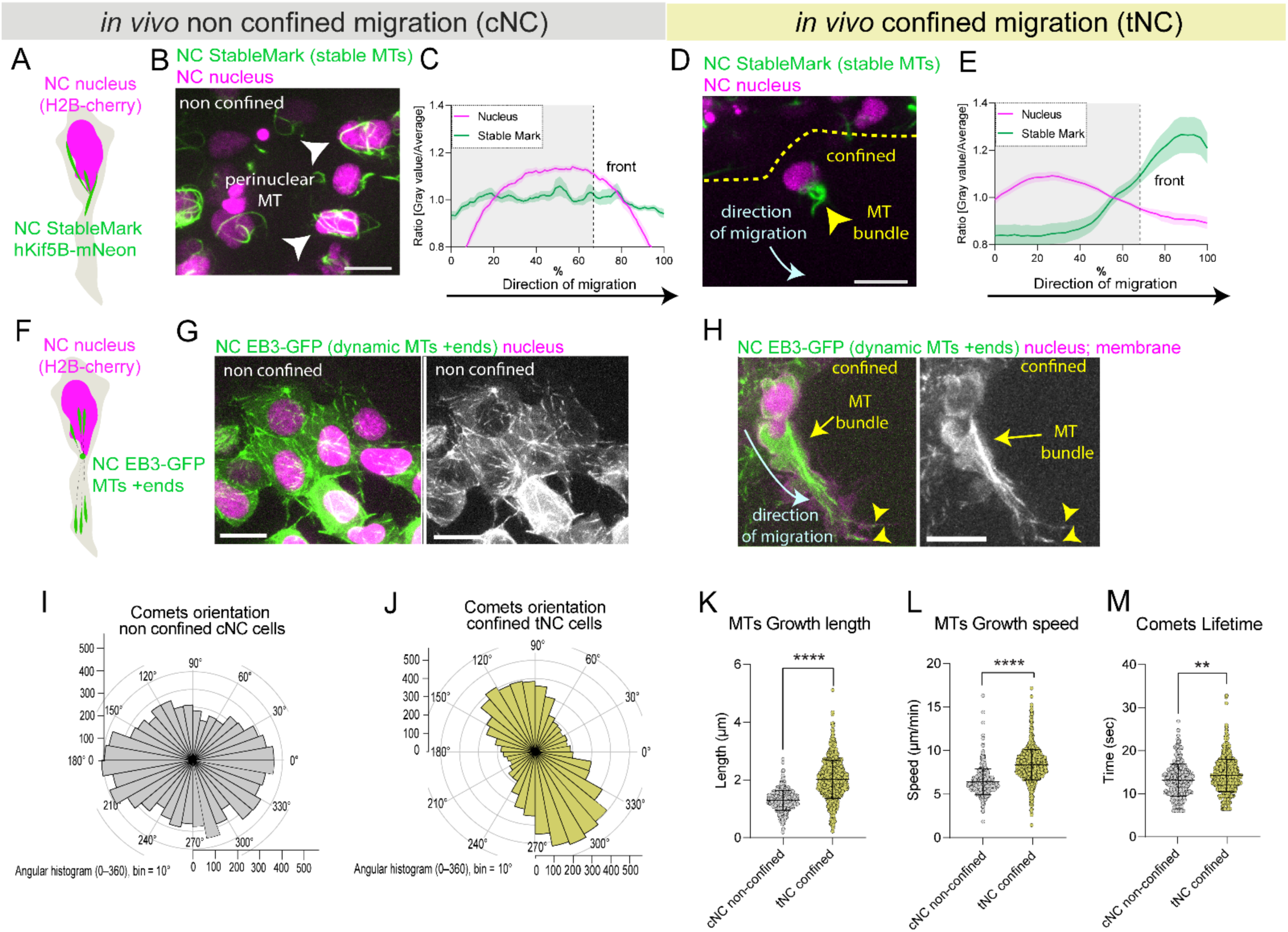
Microtubules reorganise into a frontally polarized stable bundle during confined migration *in vivo*. (A, F) Schematic representation of the transgenic zebrafish lines used in this study. Stable microtubules were visualised by expressing the rigor mutant hKif5B-NeonGreen in neural crest cells (A), while microtubule plus-end dynamics were monitored using EB3-GFP (F). In both cases, neural crest cell nuclei were labelled with H2B-mCherry. (B) Representative images of cNC cells migrating in a non-confined environment (Sox10:Kalt4-H2Bcherry / UAS:StableMark-NeonGreen; in magenta and green, respectively). Arrowheads indicate stable perinuclear microtubules. (C) Spatial distribution of stable microtubules (green) in cNC cells relative to the nucleus (magenta). Grey shadowed region represents the cell rear and centre whereas the line separate this from nucleus front. *n* = 25 cells, 4 embryos, 2 experiments. Error bars represent SEM. (D) Representative image of a tNC cell migrating in confinement (Sox10:Kalt4-H2Bcherry / UAS:StableMark-NeonGreen; magenta and green, respectively). Yellow arrowhead indicates a stable microtubule bundle at the nuclear front. Scale bar, 10 µm. (E) Spatial distribution of stable microtubules in confined tNC cells, showing enrichment at the nuclear front. Grey shadowed region represents the cell rear and centre whereas the line separate this from nucleus front. *n* = 30 cells, 5 embryos, 3 experiments. Error bars represent SEM. (G) Representative images of cNC cells (Sox10:EB3-GFP / Sox10:Kalt4-H2Bcherry / Sox10:mRFP; green and magenta respectively). Scale bar, 10 µm. (H) Representative images of a migratory tNC cell in the confined region from a Sox10:EB3-GFP / Sox10:Kalt4-H2Bcherry/ Sox10:mRFP transgenic embryo (green and magenta, respectively). Cyan arrow indicates the direction of migration and yellow arrows indicate formation of a microtubule bundle at the nuclear front while arrowheads indicate some comets reaching filopodia. Scale bar, 10 µm. (I) Angular histogram (0–360°) of EB3-GFP comet orientation in cNC cells. n = 27 cells, 6,804 comets, 6 embryos, 2 experiments. Counts range from 0 to 500. Bin size = 10°. (J) Angular histogram (0–360°) of EB3-GFP comet orientation in migratory tNC cells. n = 21 cells, 10,519 comets, 6 embryos, 3 experiments. Counts range from 0 to 500. Bin size = 10°. (K-M) EB3-GFP comet growth length (E), speed (F) and lifetime (G) in cNC non-confined cells (grey) versus tNC confined cells (yellow). cNC: *n* = 27 cells, 6,237 comets, 6 embryos, 2 experiments confined; tNC: *n* = 21 cells, 9,632 comets, 6 embryos, 3 experiments. Two tailed Mann Whitney test. **** p < 0.0001; ** p = 0.0053. Each dot represents one comet.

### Centrosomal microtubules deform the nucleus during tNC confined migration *in vivo*

To understand how microtubule reorganisation might facilitate nuclear translocation, we next examined centrosome positioning relative to nuclear deformation. In migrating neurons, extension of the leading process is accompanied by positioning of the centrosome ahead of the nucleus, which is subsequently pulled forward towards the leading edge during nucleokinesis, driving nucleus translocation (Solecki et al., 2006; Wynshaw-Boris et al., 2001; Schaar and McConnell, 2005).

During confined migration of NC cells *in vivo*, we identified the centrosome, either by live imaging of EB3-GFP (Kroll et al., 2023) (Figure 4A-C, Figure S4A, A’, Supplementary Movie 6) or by immunostaining of gamma-tubulin (Figure S4C-F). Analysis of centrosome orientation relative to the nucleus centroid revealed that the centrosome is not polarised relative to the nucleus in non-confined cranial neural crest (cNC) cells (Figure 4B, Supplementary Movie 6 cNC: arrowheads), instead appearing randomly distributed during migration (Figure 4B, D,E gray curve, F). To ask whether physical proximity of the centrosome to the nucleus might promote its deformation, we measured the distance between the nucleus centroid and the centrosome (d/R: Figure 4G)(Biedzinski et al., 2019). In cNCs, this distance is small, but constant, suggesting the centrosome lies close to the nuclear periphery (Figure 4H, I). High Z-resolution cross-section views of the cNC nucleus (Figure S4A) indeed shows that the centrosome lies above the nucleus in cNCs. Consistently, no nuclear indentation was observed in these cells (Figure 4 B,G,H gray curve, I and Figure S4A, S4B, S4C’) and we did not observe any correlation between nucleus circularity and centrosome distance from the nucleus in cNCs (Figure S4D).

**Figure 4.**
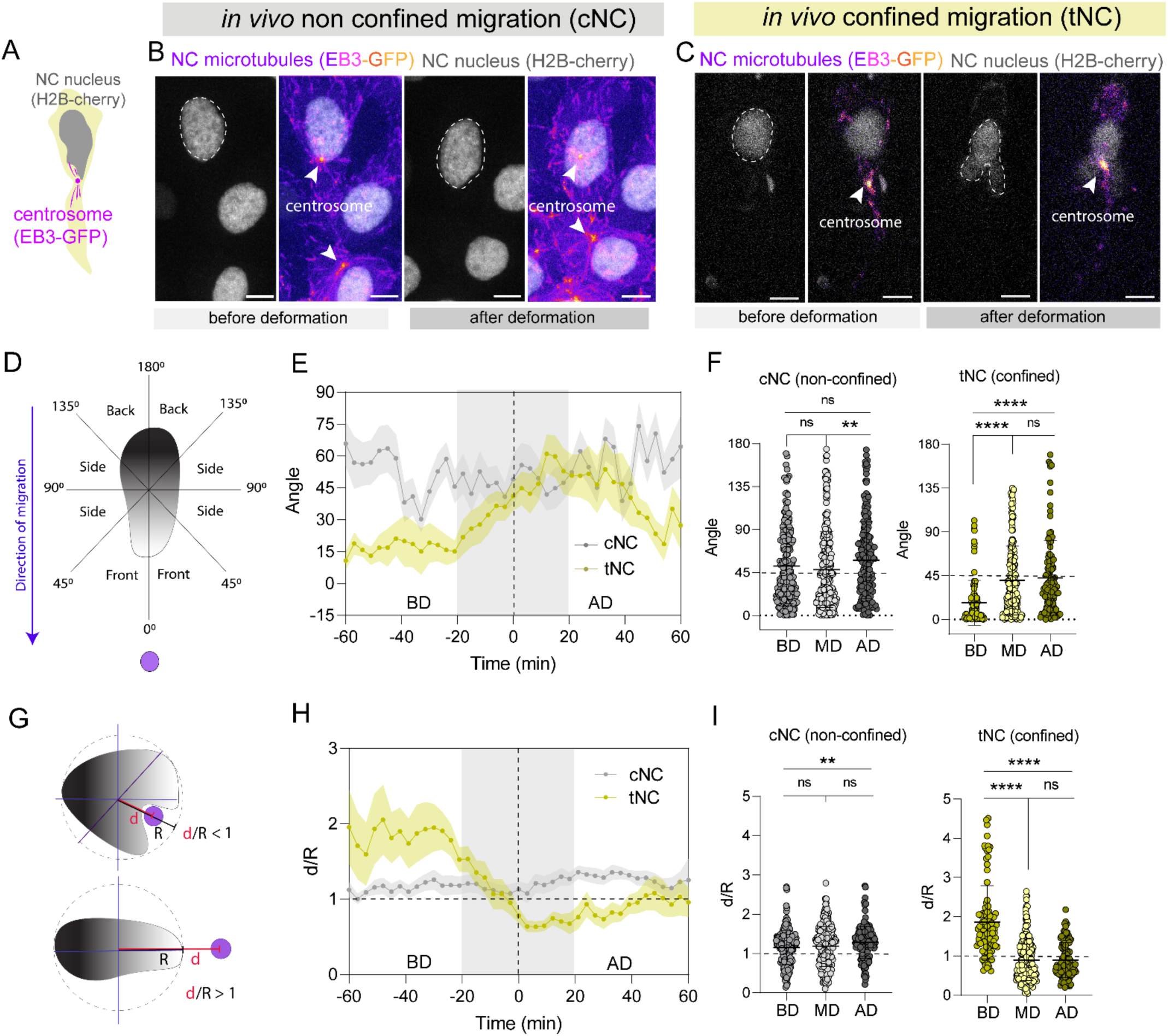
Dynamic centrosome repositioning drives transient nuclear indentation during confined tNC migration *in vivo*. (A) Schematic representation of neural crest (NC) cells expressing Sox10:EB3-GFP and H2B-Cherry to visualize centrosome orientation and nuclear dynamics during migration. (B-C) Representative images of Sox10:EB3-GFP / Sox10:Kalt4 transgenic embryos showing centrosome behaviour in non-confined cNC cells (B) and confined tNC cells (C). EB3-GFP comets are displayed in pseudocolour and nuclei in grey. Arrowheads indicate centrosome position and sites of nuclear indentation. Scale bars, 5 µm. (D) Schematic illustrating centrosome positioning analysis relative to the migration axis. Centrosome orientation was quantified from 60 min before to 60 min after the point of maximum nuclear deformation (time = 0). Angles were measured relative to the direction of migration: ∼0° indicates front polarization, ∼45° lateral positioning, and ≥90° rear positioning. (E) Mean centrosome angle relative to the direction of migration over time, aligned to maximum nuclear deformation (time = 0). In non-confined cNC cells (grey curve), centrosome positioning remained broadly lateral and showed no directional bias. In contrast, in confined tNC cells (yellow curve), the centrosome was polarized toward the front before deformation (angle < 15°) and shifted laterally (∼40°) during the period of maximum nuclear deformation and nuclear translocation (grey shaded region, −20 to +20 min). cNC: *n* = 21 cells from 7 embryos across 3 experiments. tNC: *n* = 17 cells from 9 embryos across 3 experiments. Error bars represent SEM. (F) Quantification of centrosome angle before deformation (BD; -60 to -20 min), at maximum deformation (MD; -20 to 20 min), and after deformation (AD; 20 to 60 min). Kruskal–Wallis multiple comparisons test. cNC: BD vs MD, *p* = 0.5237; MD vs AD, *p* = 0.0061; BD vs AD, *p* = 0.3262. tNC: **** *p* < 0.0001; ns *p* > 0.9999. Each dot represents one time point. cNC: *n* = 21 cells comprising 726 total time points (214 BD, 300 MD, 212 AD), from 7 embryos across 3 experiments. tNC: *n* = 17 cells comprising 444 total time points (101 BD, 240 MD, 103 AD) from 9 embryos across 3 experiments. (G) Schematic of centrosome positioning analysis during nuclear deformation. *d* is the distance between the nuclear centroid and the centrosome, and *r* is the nuclear radius (distance from centroid to nuclear envelope). A ratio *d/r* < 1 indicates that the centrosome lies within the nuclear boundary, consistent with nuclear indentation, whereas *d/r* > 1 indicates no indentation. (H) Mean centrosome indentation (d/r ratio) over time aligned to maximum nuclear deformation (time = 0). In non-confined cNC cells, the centrosome remained outside the nuclear boundary throughout the analysed time window (d/r > 1). In confined tNC cells, centrosomes remained external before deformation but transiently moved into the nuclear boundary during maximum deformation (grey shaded region, −20 to +20 min), coinciding with nuclear translocation and indentation. cNC: *n* = 21 cells from 7 embryos across 3 experiments. tNC: *n* = 17 cells from 9 embryos across 3 experiments. Error bars represent SEM. (I) Quantification of centrosome-induced nuclear indentation in cNC and tNC cells before deformation (BD; -60 to -20 min), at maximum deformation (MD; -20 to 20 min), and after deformation (AD; 20 to 60 min). MD corresponds to the time point of minimum nuclear circularity. Kruskal–Wallis multiple comparisons test for non-confined cNC cells showed no differences between BD and MD (*p* = 0.8476) nor between MD and AD (*p* = 0.0945), and limited differences (** *p* = 0.0084) between BD and AD. In contrast, for confined tNC cells showed significant differences (**** *p* < 0.0001) between BD and MD and BD and AD, whereas between MD and AD there is not significant differences (ns *p* > 0.9999). Each dot represents one time point. cNC: *n* = 21 cells comprising 726 total time points (214 BD, 300 MD, 212 AD), from 7 embryos across 3 experiments. tNC: *n* = 17 cells comprising 444 total time points (101 BD, 240 MD, 103 AD) from 9 embryos across 3 experiments.

On the other hand, in confined tNCs, centrosome positioning during nucleokinesis appears highly regulated. Indeed, the centrosome is positioned at the front of the nucleus prior to its deformation (Figure 4C, arrowhead, Figure 4D,E yellow curve, Supplementary Movie 6 tNC: arrowhead), although it remains relatively distant from the nuclear envelope (Figures 4C, G, H yellow curve and Figure S4E’). As cells undergo nuclear translocation, however, the centrosome relocates laterally to the nucleus (Figure 4D,E, yellow curve, 4F, Supplementary Movie 6 tNC: arrowhead), thus approaching the nuclear envelope and inducing a local indentation of the nucleus (Figures 4C,G,H yellow curve, 4I and Figure S4A’, S4B, S4E’, Supplementary Movie 6 tNC: arrowhead). In confined tNCs, we observed a significant positive correlation between nucleus circularity and centrosome distance from the nucleus (Figure S4F), indicating that centrosome-mediated nuclear deformation occurs during confined migration *in vivo*.

In summary, dynamic repositioning of the centrosome during confined tNC nuclear translocation suggests a functional coupling between centrosome positioning, microtubule organisation, and nuclear deformation.

### Microtubules exert pulling forces on the nucleus during confined migration *in vivo*

We find that microtubules are organised in a stable bundle in confined, migrating tNCs. Furthermore, the centrosome is polarized in front of the nucleus, and it is dynamically repositioned, possibly causing the nucleus to deform, during tNC nuclear translocation. Microtubules can generate pulling forces on the nucleus during neuronal nucleokinesis (Shu et al., 2004; Wu et al., 2018; Umeshima et al., 2019). Thus, we next asked whether a similar mechanism operates during confined neural crest migration *in vivo*. To address this, we performed IR-laser ablation experiments by tissue-specific labelling of the whole neural crest microtubule array with EMTB-3xGFP (Miller et al., 2009) and neural crest nuclei with H2B-mCherry, respectively (Alhashem et al, 2021). Subcellular IR-ablation in the region of the microtubule array in non-confined cNC cells (Figure 5A,B, Supplementary Movie 7) revealed no detectable changes in nuclear shapes (Figure 5B, C, D). This suggests that, in non-confined cells, microtubule-generated forces do not deform the nucleus.

**Figure 5.**
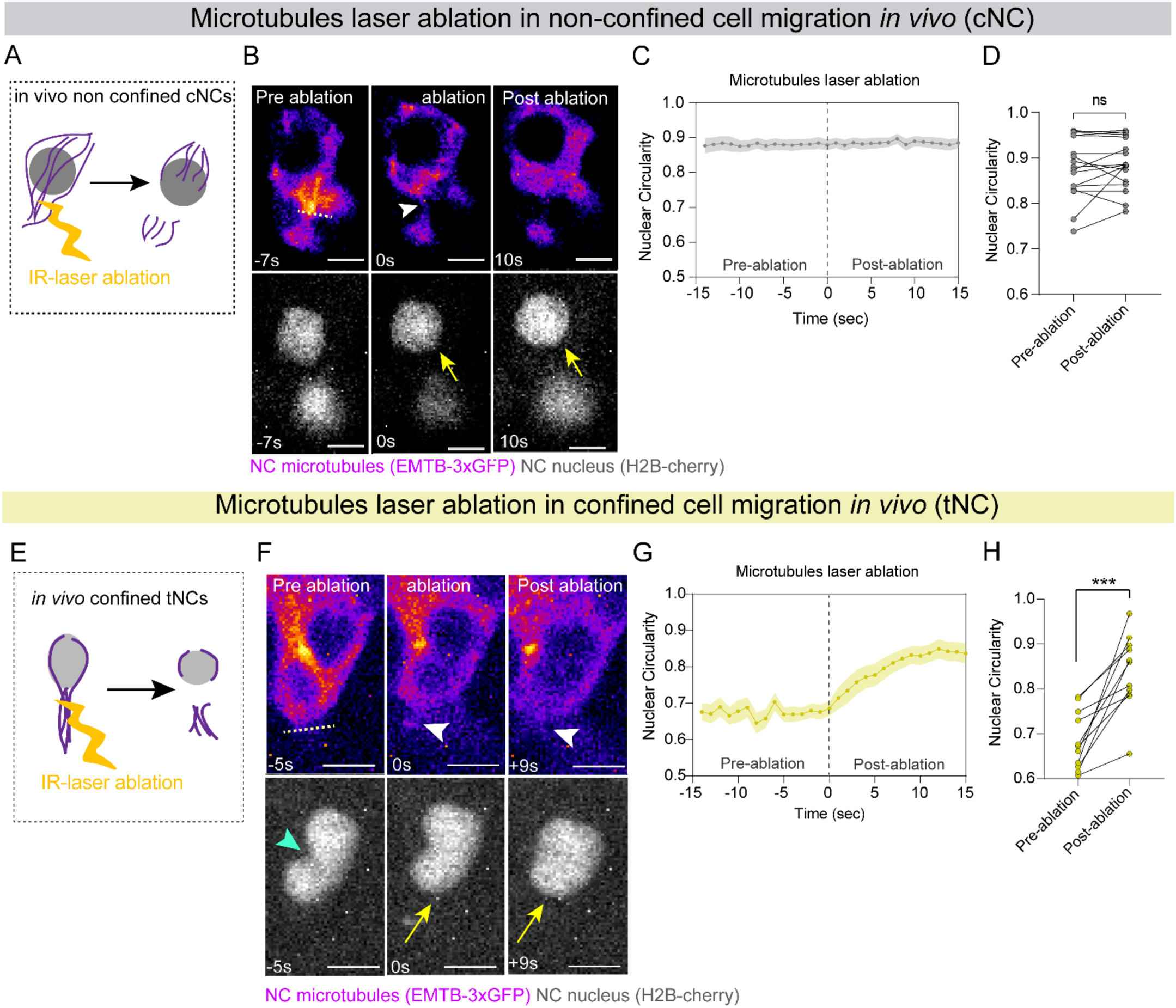
Laser ablation of microtubule bundles reveals microtubule-dependent nuclear tension during confined migration *in vivo*. (A) Schematic of a non-confined cNC cell in a live 14-somite-stage zebrafish embryo during microtubule laser ablation. Following severing, microtubule and nuclear responses were monitored. (B) Representative images of a cNC cell undergoing laser ablation (Sox10:Kalt4 / UAS:EMTB-3xGFP). Microtubules are displayed in pseudocolour and nuclei in grey. White arrowheads indicate the ablation site and yellow arrows indicate nuclear displacement. Scale bar, 5 µm. (C) Mean nuclear circularity over time aligned to the time of microtubule ablation (time = 0). cNC cells showed minimal nuclear recoil and no detectable nuclear reshaping following microtubule severing. *n* = 17 cells from 8 embryos across 2 experiments. Error bars represent SEM. (D) Quantification of nuclear circularity before and after microtubule ablation in cNC cells. No significant difference was detected (two-tailed paired *t*-test; *p* = 0.3143). *n* = 17 cells from 8 embryos across 2 experiments. (E) Schematic of a confined tNC cell in a live 21-somite-stage zebrafish embryo showing laser ablation of microtubule bundles (laser represented in yellow). Subsequent microtubule and nuclear recoil were monitored as a readout of intracellular mechanical tension. (F) Representative images of a migrating tNC cell with a deformed nucleus undergoing microtubule ablation. Microtubules are shown in pseudocolour and the nucleus in grey. White arrowheads indicate the ablation site and yellow arrows indicate nuclear recoil. Scale bar, 5 µm. (G) Mean nuclear circularity over time aligned to the point of microtubule ablation (time = 0). tNC deformed nuclei exhibited rapid recoil and changes in nuclear shape following microtubule severing, consistent with release of mechanical tension. *n* = 11 tNC cells, from 11 embryos across 9 experiments. Error bars represent SEM. (H) Quantification of nuclear circularity before and after ablation in tNC cells containing deformed nuclei. Two-tailed Wilcoxon matched-pairs signed-rank test; *** *p* = 0.0003. *n* = 11 cells from 11 embryos across 9 experiments.

In contrast, when we selectively severed the microtubule bundle positioned anterior to the nucleus in confined tNC (Figure 5E,F, S5A-C, Supplementary Movie 8), this perturbation triggered an immediate recoil of the nucleus (Figure 5F, gray scale panels). Quantification of nucleus circularity upon ablation of microtubules revealed a prompt relaxation of nuclear shape towards a more rounded configuration both in cells displaying a polarised microtubule bundle but non-deformed nuclei (Figure S5A,B, C, Supplementary Movie 9) and in cells with highly deformed nuclei (Figure 5E,G,H, Supplementary Movie 8). This rapid response indicates that microtubules exert pulling forces on the nucleus. Thus, microtubules not only transmit force but also physically deform the nucleus during confined migration *in vivo*.

Our findings support a model in which microtubules dynamically reorganise in response to confinement (Figure 3), assembling into a leading edge-oriented bundle that generates pulling forces to drive nuclear translocation (Figure 5). This confinement-dependent mechanism highlights an active role for microtubules in integrating intracellular force generation with environmental constraints to regulate nuclear positioning during NC migration.

### Dynein localises at the nuclear envelope and mediates centrosome-nucleus translocation during confined migration *in vivo*

In developing neural tissues, microtubules play a central role in nucleokinesis, by driving nucleus translocation. This process relies on dynamic interactions between the nuclear envelope and microtubule-associated motor proteins, including dynein (Wu et al., 2018; Umeshima et al., 2019; Gonçalves et al., 2020; Zhou et al., 2024). Dynein pulling forces also drive nuclear deformations in non-neuronal contexts both *in vivo* and *in vitro* including in C. Elegans zygote (Penfield et al., 2018) and immune cell differentiation (Biedzinski et al., 2019). Thus, we investigated the spatial distribution of dynein in *in vivo* migrating neural crest cells.

To this end, we examined Dynein Heavy Chain localisation in cranial neural crest (cNC) cells migrating in non-confined environments (Figure 6A) and compared it with trunk neural crest (tNC) cells migrating under confinement (Figure 6B).

**Figure 6.**
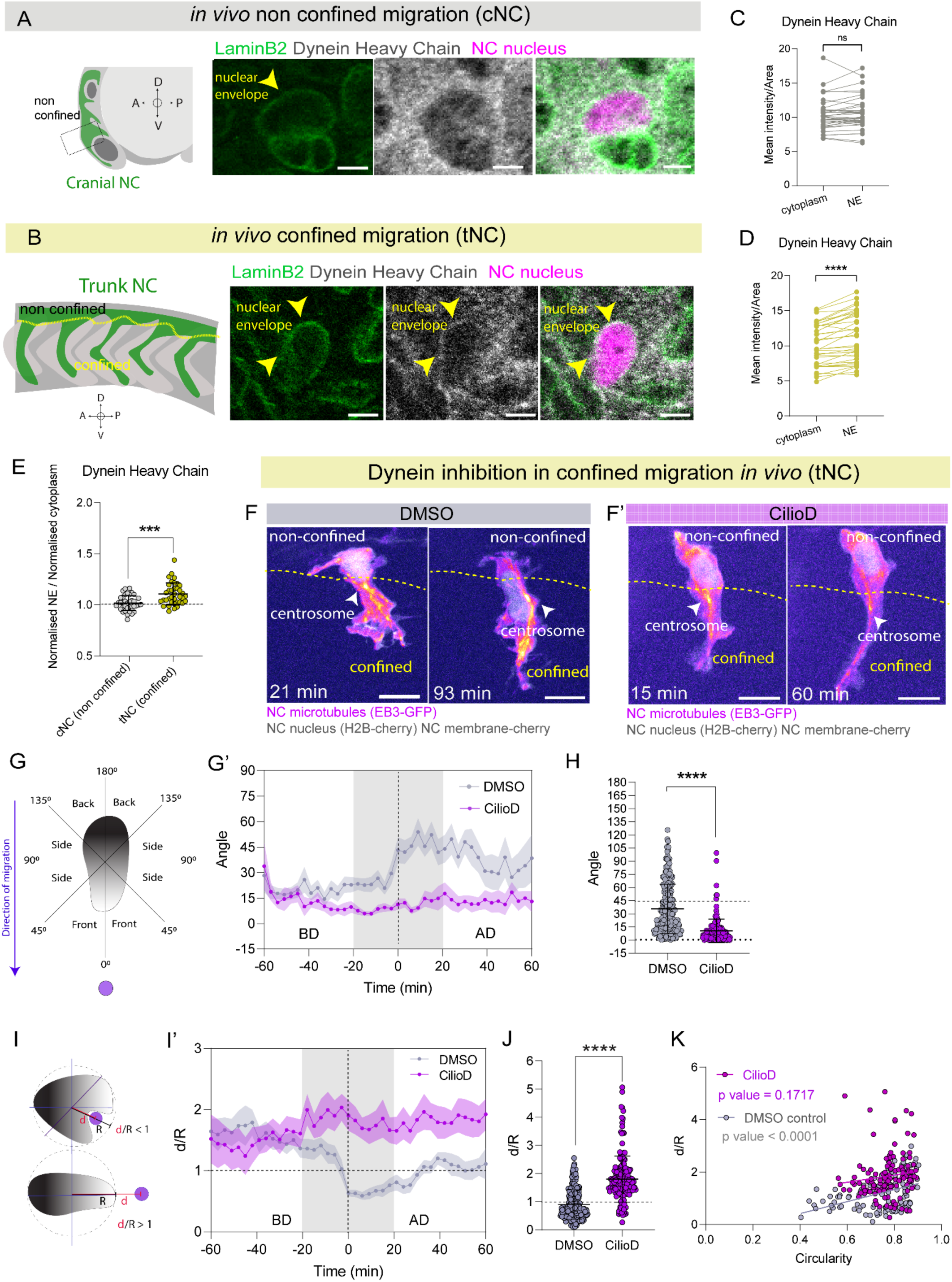
Nuclear envelope–associated dynein promotes centrosome repositioning and nuclear indentation during confined migration *in vivo*. (A) Left: schematic of cranial neural crest (cNC) migration in a 14-somite-stage zebrafish embryo within a non-confined environment. Right: representative immunofluorescence images of Lamin B2 (green) and dynein heavy chain (grey) in cNC cells (Sox10:Kalt4; nuclei in magenta). Lamin B2 outlines the nuclear envelope (arrowheads), whereas dynein remains predominantly cytoplasmic. Scale bar, 5 µm. (B) Left: schematic of the mid-trunk neural crest (tNC) region (between somites 7–10), illustrating the transition from a non-confined premigratory environment to a confined migratory environment. Right: representative immunofluorescence images of confined tNC cells stained for Lamin B2 (green) and dynein heavy chain (grey) in Sox10:Kalt4 embryos (nuclei in magenta). Arrowheads indicate dynein enrichment at the nuclear envelope co-localizing with Lamin B2. Scale bar, 5 µm. (C) Quantification of normalized dynein heavy chain intensity per area at the nuclear envelope relative to cytoplasmic regions in non-confined cNC cells. No enrichment of dynein at the nuclear envelope was detected. *n* = 37 cells from 6 embryos across 3 experiments. Two-tailed Wilcoxon matched-pairs signed-rank test; ns, *p* = 0.2246. (D) Quantification of normalized dynein heavy chain intensity per area at the nuclear envelope (NE) relative to cytoplasmic regions in confined tNC cells. Dynein is significantly enriched at the nuclear envelope. *n* = 37 cells from 9 embryos across 3 experiments. Two-tailed Wilcoxon matched-pairs signed-rank test; ****, *p* < 0.0001. (E) Ratio of normalized nuclear envelope to cytoplasmic dynein intensity in non-confined cNC and confined tNC cells. Dynein enrichment at the nuclear envelope was observed specifically in tNC cells. *n* = 74 cells in total (37 cNC and 37 tNC) from 14 embryos across 3 experiments. Mann–Whitney test; ***, *p* = 0.0002. (F-F’) Representative time series of trunk neural crest (tNC) cells from Sox10:EB3-GFP / Sox10:H2B-Cherry / Sox10:mRFP transgenic embryos transitioning from non-confined to confined regions under DMSO control and Ciliobrevin D (CilioD) treatment. In control cells (DMSO-treated), the centrosome polarizes toward the leading edge and subsequently repositions laterally to indent the nucleus during nuclear deformation. In CilioD-treated cells, the centrosome remains frontally positioned and fails to reposition during deformation. Arrowheads indicate centrosome position. Scale bar, 10 µm. (G) Schematic of centrosome positioning analysis relative to the migration axis over time, quantified from 60 min before to 60 min after maximum nuclear deformation (time = 0). Angles indicate centrosome orientation relative to the migration direction: ∼0° corresponds to front polarization, ∼45° to lateral positioning, and ≥90° to rear positioning. (G’) Mean centrosome angle over time aligned to maximum nuclear deformation (time = 0). In control cells (grey curve), the centrosome is polarized toward the leading edge before deformation and subsequently shifts laterally during the deformation period (grey shaded region, −20 to +20 min), consistent with centrosome repositioning. In Ciliobrevin D–treated cells (yellow curve), the centrosome remains frontally polarized and fails to reposition. DMSO: *n* = 20 cells from 11 embryos across 5 experiments; CilioD: *n* = 14 cells from 8 embryos across 5 experiments. Error bars represent SEM. (H) Quantification of centrosome angle at the point of maximum nuclear deformation. Two-tailed Mann–Whitney test; ****, *p* < 0.0001. DMSO: *n* = 20 cells, 291 time points from 11 embryos across 5 experiments; CilioD: *n* = 14 cells, 177 time points from 8 embryos across 5 experiments. Each dot represents one time point. (I) Schematic of centrosome indentation analysis during nuclear deformation. *d* represents the distance between the centrosome and the nuclear centroid, and *r* represents the nuclear radius (distance from the centroid to the nuclear envelope). A ratio d/r < 1 indicates centrosome penetration into the nuclear boundary consistent with nuclear indentation, whereas d/r > 1 indicates absence of indentation. (I’) Mean centrosome indentation (d/r ratio) over time aligned to maximum nuclear deformation (time = 0). In control cells (DMSO-treated, grey curve), the centrosome remains outside the nuclear boundary before deformation but transiently moves within the nuclear boundary during the deformation period (grey shaded region, −20 to +20 min), coinciding with nuclear translocation and indentation. In Ciliobrevin D–treated cells (yellow curve), the centrosome remains external to the nucleus throughout the analysed time window. DMSO: *n* = 20 cells from 11 embryos across 5 experiments; CilioD: *n* = 14 cells from 8 embryos across 5 experiments. Error bars represent SEM. (J) Quantification of nuclear indentation at the point of maximum nuclear deformation. Two-tailed Mann–Whitney test; ****, *p* < 0.0001. DMSO: *n* = 20 cells, 291 time points from 11 embryos across 5 experiments; CilioD: *n* = 14 cells, 177 time points from 8 embryos across 5 experiments. Each dot represents one time point. (K) Linear regression analysis of centrosome indentation (d/r ratio) versus nuclear circularity over time in DMSO- and CilioD-treated cells. In control cells, increased nuclear circularity (≈1) correlates with greater centrosome distance from the nuclear centroid, whereas decreased nuclear circularity (greater nuclear deformation) correlates with centrosome proximity and indentation (d/r < 1). This relationship was significant (*p* < 0.0001 from *n* = 10 cells followed, 119 timepoints). In contrast, under Ciliobrevin D treatment, this correlation was lost and centrosomes remained distant from the nucleus throughout the analysed period (*p* = 0.1717 from *n* = 14 cells followed, 126 timepoints). Each dot represents one time point.

In non-confined cNC cells *in vivo*, dynein was not enriched at the nuclear envelope (Figure 6A,C), displaying instead a predominantly cytoplasmic distribution when compared to Lamin B2 (Figure S6A). In contrast, confined tNC cells exhibited a clear enrichment of dynein at the nuclear envelope (Figure 6B, arrowheads). Quantitative analysis revealed that nuclear envelope-localised Dynein was significantly higher than Dynein levels in the surrounding cytoplasm (Figure 6D). A similar enrichment was observed for the nuclear lamina component LaminB2 (Figure S6B), which is expressed in tNCs (Hakkinen et al., 2025), indicating a robust recruitment of dynein to the nuclear periphery under confinement.

Direct comparison between the two conditions revealed a significant increase in perinuclear dynein localisation in confined tNC cells relative to non-confined cNC cells (Figure 6E). Together, these results indicate that Dynein is recruited to the nuclear envelope under confinement. Mechanical constraints might promote the assembly of a dynein-mediated microtubule–nucleus coupling system, potentially enabling the transmission of pulling forces required for nuclear deformation and translocation during confined NC migration.

To investigate the functional role of Dynein in tNC centrosome positioning, we next perturbed its motor activity using Ciliobrevin D (CilioD), a highly specific inhibitor of the AAA+ ATPase activity of cytoplasmic dynein (Biedzinski et al., 2019; Roosien et al., 2015) (Figure 6F-F’, Supplementary Movie 10). Using EB3-GFP to identify the centrosome, H2B-mCherry to mark neural crest nuclei and membrane-targeted RFP to visualise the neural crest plasma membrane, we tracked centrosome positioning relative to the nucleus centroid. Control cells exhibited dynamic centrosome repositioning during nucleus translocation (Figure 6F, G-G’ grey curve, Supplementary Movie 10, DMSO: arrowhead). In DMSO-treated embryos, the centrosome initially polarised at the front of the nucleus (Figure 6F), before repositioning laterally along the nuclear envelope. This lateral movement was accompanied by pronounced nuclear indentation and deformation, suggesting an active mechanical coupling between the centrosome and the nucleus (Figure 6F,6G-G’,6I-I’ grey curves).

In contrast, inhibition of Dynein with 15 μm CilioD at the onset of tNC migration (18 hpf) resulted in a striking defect in centrosome positioning (Figure 6F’, Supplementary Movie 10, CilioD: arrowhead). In treated embryos, tNC cells failed to reposition the centrosome, which remained persistently positioned at the front of the nucleus (Figure 6G-G’ purple curve). The ratio between the distance of the centrosome and nucleus centroid radius (d/R, Figure 6I) remained significantly higher in CilioD treated embryos (Figure 6I-J purple curve, Figure S6D). Correlation analysis between d/R and nucleus circularity indeed showed that the centrosome no longer indents the nucleus upon inhibition of Dynein motors (Figure 6K, Supplementary Movie 10, CilioD: arrowhead). To further quantify this effect, we analysed centrosome positioning in fixed embryos stained for γ-tubulin (Figure S6E-E’). In CilioD–treated embryos, the centrosome was frequently displaced away from the nuclear envelope, indicating a loss of centrosome–nucleus coupling (Figure S6F). Similarly to our live EB3-GFP data, CilioD treatment resulted in a loss of correlation between centrosome distance from the nucleus and nucleus circularity (Figure S6G).

Together, these results demonstrate that dynein activity is essential for centrosome repositioning and for maintaining its physical association with the nucleus.

### Dynein controls nucleokinesis in confined tNC cells *in vivo*

To ask directly whether Dynein drives nuclear translocation during confined migration *in vivo,* we inhibited cytoplasmic Dynein using two complementary approaches: pharmacological inhibition with Ciliobrevin D (Fig. 7A–B, Supplementary Movie 11) and neural crest-specific disruption of dynein activity via overexpression of Dynamitin, which uncouples the dynactin complex from Dynein (Melkonian et al., 2007) (Fig. 7L-M, Supplementary Movie 12).

**Figure 7.**
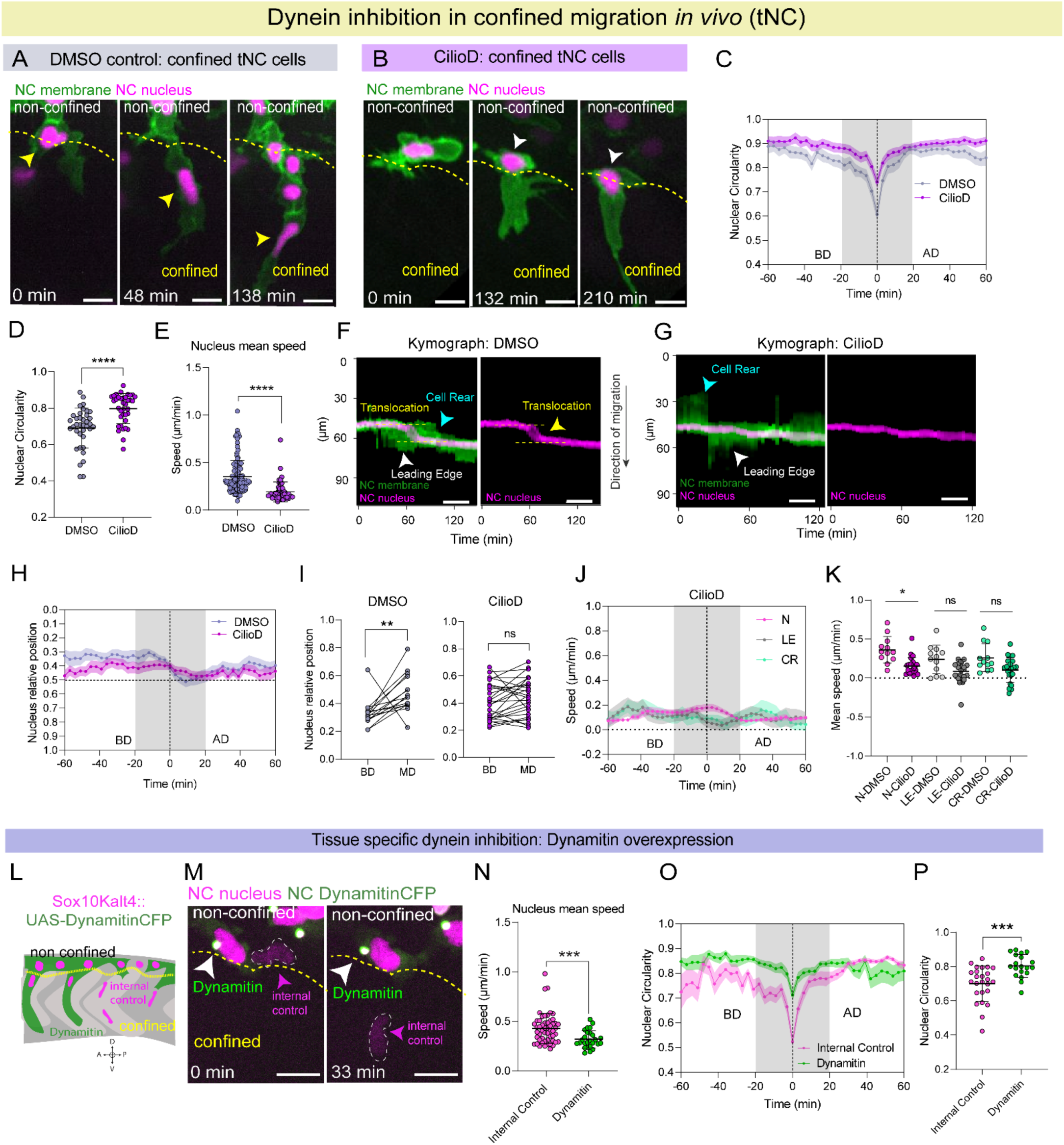
Dynein drives nuclear translocation and deformation during confined migration *in vivo*. (A-B) Representative images of tNC cells in embryos treated with DMSO (A) and CilioD (B). . Yellow arrowheads indicate nuclear translocation and deformation, whereas white arrowheads indicate failure of these processes. Scale bar, 10 µm. (C) Mean nuclear circularity over time aligned to the point of maximum nuclear deformation (time = 0), spanning 60 min before and after alignment in DMSO- and Ciliobrevin D–treated cells. DMSO: *n* = 34 cells from 10 embryos across 4 experiments; CilioD: *n* = 38 cells from 11 embryos across 4 experiments. Error bars represent SEM. (D) Quantification of mean nuclear circularity during the nuclear translocation window (−20 to +20 min relative to maximum nuclear deformation) in DMSO- and Ciliobrevin D–treated embryos. Each dot represents one cell. Two-tailed Mann–Whitney test; ****, *p* < 0.0001. DMSO: *n* = 34 cells from 10 embryos across 4 experiments; CilioD: *n* = 38 cells from 11 embryos across 4 experiments. (E) Quantification of mean nuclear speed in DMSO- and Ciliobrevin D–treated cells. Dynein inhibition significantly reduced nuclear speed. DMSO: *n* = 34 cells, 121 tracks; CilioD: *n* = 38 cells, 52 tracks. Two-tailed Mann–Whitney test; ****, *p* < 0.0001. Each dot represents one track. (F-G) Representative kymographs of membrane and nuclear intensity profiles aligned to the nuclear centroid along the migration axis in DMSO (F) and Ciliobrevin D (G) conditions. Cell rear dynamics are indicated by cyan arrowheads and leading-edge dynamics by white arrowheads. Nuclear translocation events are indicated by yellow arrowheads. Nuclear translocation was not detected following dynein inhibition. Scale bar, 10 µm. (H) Relative nuclear position within the cell over time in DMSO-treated (grey) and Ciliobrevin D–treated (purple) embryos, aligned to maximum nuclear deformation (time = 0). Values approaching 0 indicate rear localization, values near 0.5 indicate central positioning, and values approaching 1 indicate localization toward the leading edge. The grey shaded region indicates the nuclear translocation period. BD, before deformation; AD, after deformation. DMSO: *n* = 15 cells from 4 embryos across 3 experiments; CilioD: *n* = 27 cells from 6 embryos across 3 experiments. Error bars represent SEM. (I) Quantification of relative nuclear position before deformation (BD) and at maximum nuclear deformation (MD) in DMSO and Ciliobrevin D conditions. Nuclear repositioning was detected in control cells but not following dynein inhibition. Two-tailed Wilcoxon matched-pairs signed-rank test; DMSO: ** *p* = 0.0067; CilioD: ns, *p* = 0.0731. DMSO: *n* = 15 cells from 4 embryos across 3 experiments; CilioD: *n* = 27 cells from 6 embryos across 3 experiments. (J) Mean nuclear speed (N, magenta), leading-edge speed (LE, grey), and cell rear speed (CR, cyan) over time in Ciliobrevin D–treated embryos aligned to maximum nuclear deformation (time = 0). Data span 60 min before and after alignment. The grey shaded region indicates the analysis window (−20 to +20 min). Dynein inhibition markedly reduced nuclear speed. CilioD: *n* = 27 cells from 6 embryos across 3 experiments. (K) Statistical comparison of mean nuclear, leading-edge, and cell rear speeds during the analysis window (−20 to +20 min relative to maximum nuclear deformation) between DMSO- and Ciliobrevin D–treated embryos. Kruskal–Wallis multiple comparisons test revealed a significant reduction in nuclear speed following dynein inhibition (*p* = 0.0151), whereas leading-edge (*p* = 0.1986) and cell rear (*p* = 0.4453) dynamics were not significantly affected. DMSO: *n* = 12 cells from 4 embryos across 3 experiments; CilioD: *n* = 23 cells from 6 embryos across 3 experiments. (L) Schematic of tissue-specific dynein inhibition in tNC cells transitioning from non-confined to confined environments using Sox10:Kalt4 / UAS:Dynamitin-CFP, induced by tamoxifen. Dynamitin-positive cells display green cytoplasm and magenta nuclei, whereas neighbouring Dynamitin-negative cells (magenta nuclei only) serve as internal controls. (M) Representative images of confined tNC cells expressing Dynamitin-CFP. White arrowheads indicate Dynamitin-positive cells (green cytoplasm with a centrosomal signal), whereas magenta arrowheads indicate neighbouring internal controls. Scale bar, 10 µm. (N) Quantification of mean nuclear speed in control and Dynamitin-expressing cells. Dynein inhibition significantly reduced nuclear speed. Internal control: *n* = 25 cells, 53 tracks; Dynamitin: *n* = 19 cells, 26 tracks. Two-tailed Mann–Whitney test; ***, *p* = 0.0006. (O) Mean nuclear circularity over time aligned to the point of maximum nuclear deformation (time = 0) in control and Dynamitin-expressing confined tNC cells. Internal control: *n* = 25 cells from 8 embryos across 2 experiments; Dynamitin: *n* = 19 cells from 7 embryos across 2 experiments. Error bars represent SEM. (P) Quantification of mean nuclear circularity during the nuclear translocation window (−20 to +20 min relative to maximum nuclear deformation) comparing internal controls and Dynamitin-positive cells. Each dot represents one cell. Two-tailed unpaired *t*-test; ***, *p* = 0.0004. Internal control: *n* = 25 cells from 8 embryos across 2 experiments; Dynamitin: *n* = 19 cells from 7 embryos across 2 experiments.

Disruption of Dynein motor activity led to a severe impairment in nuclear translocation and in overall confined tNC migration (Fig. 7A-B,7L–M, Supplementary Movie 11 and 12). Strikingly, Ciliobrevin D-treated nuclei did not undergo deformation (Fig. 7C-D) and instead remained positioned at the rear of the cell (Figure 7B: arrowheads, Supplementary Movie 11: compare yellow arrows). As a consequence, cells were unable to efficiently translocate their nucleus from the premigratory (non-confined) region into the confined migratory zone (Figure 7A,B, Supplementary Movie 11: compare yellow arrows). A similar phenotype was observed upon neural-crest specific Dynamitin overexpression: while neighbouring Dynamitin negative cells (internal controls; Fig. 7M, magenta arrows, Supplementary Movie 12: magenta arrow) successfully entered confined regions, undergoing nucleus translocation and deformation, Dynamitin-expressing cells (Fig. 7M, white arrowheads, Supplementary Movie 12: white arrowhead) failed to do so. Measurement of nucleus speed (Figure 7E,7N) and nucleus circularity (Figure 7C-D, 7O-P) reveals that tNC nucleokinesis and nuclear deformation is impaired upon disruption of Dynein function. To quantify membrane dynamics relative to nucleus position (Figure 7F, G), we carried out membrane and nucleus segmentation and 1D kinematic analysis of the leading edge and cell rear upon Ciliobrevin D treatment. Intracellular nucleus translocation failed in Dynein inhibited cells, and the nucleus remained close to the cell centre (Figure 7H-I). We observed that the speed of nucleus translocation was strongly reduced upon Dynein inhibition (Figure 7J-K, S7C). Leading edge speed before nuclear translocation was significantly affected by Dynein inhibition (Figure 7J, S7A,S7D) whilst cell rear retraction after deformation remained unchanged (Figure 7J, S7B, S7E). Cross correlation analysis of nucleus against leading edge or cell rear speeds showed these were not correlated upon Ciliobrevin D treatment in tNCs (Figure S7F).

To determine whether the requirement for Dynein for nucleus translocation is specific to confined migration, we performed equivalent perturbations in cNC cells, which migrate in non-confined environments. Using both pharmacological inhibition (Fig. S7G-H, Supplementary Movie 13) and neural-crest specific Dynamitin overexpression (Fig. S7P-S, Supplementary Movie 14, compare magenta arrows with white arrowheads), we found that dynein disruption had no detectable effect on nuclear shape changes(Fig. S7I-I’), nuclear positioning (Fig. S7J-K), or nucleus, leading edge or cell rear migration speed (Fig. S7L,M). Cross correlation analysis of nucleus against cell rear or leading edge speed indicates these remain highly correlated in Dynein motor inhibited cells as well as in the control (Figure S7N, O), thus maintaining coordination between nucleus and cell membrane movement.

Together, these results demonstrate that Dynein is essential for nucleokinesis of tNCs during *in vivo* migration through confinement, where it enables nuclear deformation, nuclear translocation and tNC developmental migration. By contrast, Dynein is dispensable for nuclear dynamics during non-confined migration of cNCs, highlighting a novel, confinement-dependent mechanism in which Dynein-mediated pulling forces are required to overcome physical constraints and drive nuclear translocation.

## Discussion

We discovered that during developmental migration through confined tissue environments, tNC cells undergo a Dynein-dependent nucleokinesis process, in which intracellular nuclear translocation is closely coordinated, but mechanistically uncoupled, from actomyosin cortex and plasma membrane dynamics. Microtubule motors are known to drive nuclear translocation across the evolutionary tree, spanning from fungi (Morris et al., 1998), to C. Elegans (Fridolfson and Starr, 2010), to mammalian myotubes (Wilson and Holzbaur, 2012, Wilson and Holzbaur, 2015) and mammalian central nervous system development (Tsai et al., 2005). To our knowledge, dynein-dependent nuclear translocation has not been yet reported as a means to achieve migration through confinement.

Our quantitative analysis of nucleus versus plasma membrane kinematics revealed that at the time of maximal nuclear deformation, the movement of the leading edge and cell rear appear independent of nucleus translocation (Figure 1B’, 1J-L’, S1G-G’). We find that leading edge protrusion precedes nuclear translocation, which is followed by a delayed cell rear contraction. Live imaging of Myosin-light-chain-GFP in confined tNCs reveals that a small pool of Myosin II transiently localises at the cell front, followed by a marked accumulation at the cell rear (Figure 2B). Perturbation of actomyosin contractility by pharmacological inhibition of ROCK (Figure 2D) results in elongated cells characterised by slightly delayed cell rear retraction and mildly impaired leading edge advancement. These phenotypes are compatible with Myosin-II dependent regulation of actin retrograde flow at the leading edge (Medeiros et al., 2006) and of trailing edge retraction (Garrido-Casado et al., 2021). ROCK inhibition neither affects nucleus shape changes nor the speed of nucleus translocation, indicating that actomyosin is not required for these processes. *In vitro* approaches (Liu et al., 2015, Bergert et al., 2012) illuminated a role for actomyosin contractility for migration through confinement. Highly confined cells can use their nucleus as a proprioceptive device to measure their surroundings (Renkawitz et al., 2019) and respond to nucleus deformations by upregulating actomyosin contractility, which facilitates rapid migration through confinement (Lomakin et al., 2020, Venturini et al 2020). Nuclear translocation in confined cell migration often requires actomyosin pushing forces at the cell rear (Thomas et al., 2015, Keys et al., 2024, Ju et al, 2024) or perinuclear Arp2/3 mediated actin polymerization (Thiam et al., 2016) to drive nuclear passage through confinement.

By contrast, we observe that, despite extreme nucleus deformations, in tNCs contractility appears dispensable for nucleokinesis. This discrepancy might be linked to the likely softer composition of the tNCs nuclear envelope (Hakkinen et al., 2025), which might impair nuclear mechanosensing (Lomakin et al., 2020), to the soft nature of the embryonic tissue environment traversed by neural crest cells (Marchant et al., 2022) or to activation of microtubule-dependent adaptive responses to confinement (Li et al., 2023 Ju et al., 2024, Hunter et al., 2025).

Indeed, our data indicate that in confined tNCs, but not in non-confined cNCs, intracellular microtubule-dependent *pulling f*orces drive nucleokinesis *in vivo*. The mode of migration of tNCs under confinement appears qualitatively akin to mammalian cerebellar granule neurons migration (Umeshima et al., 2018). Tissue-specific *in vivo* live imaging of stable and dynamic microtubule subpopulations reveals that microtubules reorganise from a perinuclear mesh into a leading edge-oriented bundle (Figure 3), which may act as a structural “highway” guiding nuclear translocation under confinement. The centrosome, which acts as the main microtubule organising centre in tNC cells, indents and deforms the nuclear periphery during translocation (Figure 4). IR-laser ablation of the microtubule bundle results in restoration of nuclear circularity in confined, but not in non-confined, NCs (Figure 5). Thus, we discover that microtubule-dependent pulling forces drive nuclear movement in confined tNCs. A pool of dynein-heavy-chain localises at the nuclear envelope in confined tNCs, and both centrosome repositioning around the nucleus and nucleokinesis are mediated by dynein motors (Figure 6 and 7). Dynein motor activity also regulates leading edge speed in confined tNCs (Figure S7D). This phenotype is reminiscent of neuronal growth cone extension, where loss of dynein function leads to defective trafficking of microtubules to growth cone filopodia (Myers et al., 2006) and to impaired pushing forces at the leading edge (Roosien et al., 2014).

In postmitotic migrating neurons, the LINC complex components Nesprin2 and Sun2/Sun1 anchor dynein to the nuclear periphery (Zhang et al., 2009; Zhou et al., 2024). tNC are actively proliferating precursors of the peripheral nervous system (Prendergast and Raible, 2014; Le Douarin and Smith, 2003), and cell-cycle regulation is required for correct collective migration (Alhashem et al., 2022). As nuclear anchoring of dynein can also occur independently of the LINC complex via nuclear pore complex components during interkinetic nuclear migration of cycling neural progenitors (Hu et al., 2013), future work will elucidate how dynein might mediate force transmission to the nucleus in tNCs.

By contrast, the non-confined counterpart of tNCs, cranial NCs, do not require Dynein motors for their collective migration, and both membrane and nucleus shapes and movement are tightly coupled in these cells (Figure S7). Dynein does not localise at the nuclear envelope in non-confined cranial NCs (Figure 6), suggesting that a switch to dynein-dependent nucleocytoplasmic coupling may occur specifically in confined NCs.

Quantitative analysis of microtubule growth dynamics reveals that tNC are characterized by fast growing, highly oriented microtubules at the cell leading edge, whilst microtubule growth speed, growth length and co-orientation is considerably lower in non-confined cNCs (Figure 3). Microtubules resist compressive loads and can be stabilized in the cytoplasm in response to physical confinement either via acetylation (Xu et al., 2017) or via recruitment of CLASPs to the microtubule lattice (Li et al. 2023, Ju et al. 2024). Thus, under confinement tNCs microtubules may adaptively stabilize, thus favouring a switch to a nucleokinesis mode of migration by facilitating dynein force-generation (Estrem et al., 2017).

An alternative scenario may entail transcriptional upregulation of a neuronal-like invasion program as a response to physical confinement (Hunter et al., 2025). Neural crest are the multipotent precursors of neurons and glia of the peripheral nervous system (Le Douarin and Smith, 2003). Neuronal-specific tubulins are transcriptionally upregulated in migrating chick NCs (Echeverria et al., 2026). Zebrafish and human invasive melanomas, which are NC-derived and upregulate neural crest states during metastasis (Kaufman et al., 2016) can switch to a microtubule-dependent neuronal invasion programme in confinement (Hunter et al., 2025). Thus, neural crest may similarly switch to a neuronal-like nucleokinesis programme as they encounter confined tissue environments in development. What could be the advantage of using nucleokinesis as confined migration modality? Both developing CNS neurons and neural crest cells are likely endowed with soft nuclei, as they do not express LaminA/C (Hakkinen et al., 2025, Chen et al., 2019, Amini et al., 2022). Moreover, these cell types traverse soft, but often densely packed, tissue environments during their development (Iwashita et al., 2014, Amini et al., 2022, Marchant et al., 2022). Thus, their soft nucleus might be efficiently transported by motors as a deformable cargo, enabling negotiation of tissue confinement unhindered by nuclear stiffness. Microtubule-dependent mechanisms of confined migration might also underlie invasiveness of brain cancers, such as gliomas, specialised in navigating through soft brain tissues (Cuddapah et al., 2014). In summary, our work provides insight in how an evolutionarily successful, multipotent cell type, the neural crest, adapts its cytoskeletal organisation to challenging extracellular environments, ensuring robust developmental migration in the early embryo.

## Materials and Methods

### Molecular cloning

To enable expression of transgenes under the Gal4/UAS system and facilitate identification of transgenic zebrafish embryos, the 14xUAS-E1b:BGi-nls-emGFP (plasmid kindly provided by Harol Burgess; Bergeron et al., 2015), was modified to include a cmlc2-eGFP expression cassette, which drives eGFP expression in the heart for screening purposes. The cmlc2-eGFP cassette (cardiac myosin light chain 2 fused to eGFP) was amplified from pDestTol2CG2 (Kawakami et al., 2004; Urasaki et al., 2006) using the primers 5′-CATCATGGTACCAAAGCTTAAATCAGTTGT-3′ and 5′-ATGATGGGTACCTTACTTGTACAGCTCGTC-3′. The resulting fragment was inserted into the KpnI site of the 14xUAS-E1b:BGi-nls-emGFP vector using T4 DNA ligase (Promega), generating a modified backbone containing a cmlc2-driven eGFP reporter for heart-specific fluorescence screening.

This plasmid was subsequently used as the backbone for all constructs described below.

The nls-emGFP cassette was removed by digestion with EcoRV-HF (New England Biolabs, 10151158) and SpeI-HF (New England Biolabs, 10192521). The digested DNA was resolved on a 1% agarose gel in TAE buffer (∼100 V), and the ∼7,115 bp UAS backbone fragment (containing 14xUAS-E1b:BGi cmlc2-eGFP) was excised and purified using the QIAquick Gel Extraction Kit (Qiagen, 169012830).

EMTB-3xGFP and StableMark-NeonGreen constructs were subcloned into this UAS backbone to enable targeted expression in neural crest cells.

For the UAS:StableMark-NeonGreen construct, the insert was amplified from the plasmid pB80-hKif5b(1–560)-rigor-2×mNeonGreen (Addgene plasmid # 174649 ; http://n2t.net/addgene:174649 ; RRID:Addgene_174649; Jansen et al., 2023). PCR amplification was performed using the forward primer 5′-CTGGGCAACGTGGAATTCGATATCATGGCGGACCTGGCCGAG-3′ and the reverse primer 5′-GGTGGCTGAGACTTAATTACTAGTTTACTTGTACAGCTCGTCCATGCC-3′, which introduced EcoRV and SpeI restriction sites. Reactions were carried out using CloneAmp HiFi PCR Premix (Takara, 2209559A) under the following cycling conditions: 98 °C for 30 s; 30 cycles of 98 °C for 10 s, 65 °C for 30 s, and 72 °C for 1 min; followed by a final extension at 72 °C for 2 min.

PCR products were resolved on a 1% agarose/TAE gel (∼100 V), and the ∼3,157 bp fragment corresponding to the StableMark insert was excised and purified. The insert and UAS backbone (both digested with EcoRV and SpeI) were assembled using the NEBuilder HiFi DNA Assembly Cloning Kit (New England Biolabs, 10173994).

The resulting plasmid was transformed into competent cells, and single colonies were selected and cultured overnight in LB broth containing ampicillin. Plasmid DNA was extracted using the Monarch Plasmid Miniprep Kit (New England Biolabs, 10132561). Positive clones were identified by PCR screening, with correct constructs yielding a ∼3,157 bp band following agarose gel electrophoresis. Confirmed clones were further expanded, and plasmid DNA was purified using the HiSpeed Plasmid Midi Kit (Qiagen, 12643).

### Generation of Transgenic Zebrafish

#### UAS-StableMark-NeonGreen

To generate the UAS:StableMark-NeonGreen transgenic line, 10 pg of pTol1-UAS:hKif5b(1–560)-rigor-2×mNeonGreen cmlc2-eGFP plasmid DNA was co-injected with 80 pg of Tol1 transposase mRNA into one-cell stage embryos. F1 founders were identified based on green fluorescence in the heart.

#### UAS-EMTB-3XGFP

To generate the UAS:EMTB-3xGFP construct, the EMTB-3xGFP insert (Addgene plasmid # 26741 ; http://n2t.net/addgene:26741 ; RRID:Addgene_26741; Miller et al., 2009) was amplified by PCR using the forward primer 5′-CTGGGCAACGTGGAATTCGATATCATGGAGCAGAAGCTCATCTCAGAA-3′ and the reverse primer 5′-GGTGGCTGAGACTTAATTACTAGTTTACTTGTACAGCTCGTCCATGCC-3′. The amplified fragment was then ligated into the linearised UAS backbone, as described above, using T4 DNA ligase (Promega) and subsequently transformed into competent bacterial cells.

Transgenic lines were generated by co-injecting 10 pg of pTol1-UAS:EMTB-3xGFP cmlc2-eGFP plasmid DNA with 80 pg of Tol1 transposase mRNA into embryos at the one-cell stage. F1 founders were identified by screening for green fluorescence in the heart.

#### UAS-Dynamitin-CFP

The plasmid p5UAS-3xCitrineDynamitinGI was kindly provided by Dr. Reinhard Köster (Theisen et al., 2020). The Dynamitin-CFP fragment was subcloned from this plasmid into the 14xUAS-E1b:BGi cmlc2-eGFP backbone to enable Tol1-mediated integration and heart-specific GFP screening. The Dynamitin-CFP insert was amplified by PCR using the following primers: forward 5′-CTGGGCAACGTGGAATTCGATATCATGGTGAGCAAGGGCGAGGAGC-3′ and reverse 5′-GGTGGCTGAGACTTAATTACTAGTCTACTTGTTGAGTTTCTTCATC-3′. PCR products were subsequently ligated into the linearised backbone using T4 DNA ligase (Promega) and transformed into competent bacterial cells. To generate the UAS:Dynamitin-CFP transgenic line, 15 pg of pTol1-UAS:Dynamitin-CFP cmlc2-eGFP plasmid DNA was co-injected with 80 pg of Tol1 transposase mRNA into embryos at the one-cell stage. F1 founders were identified by screening for green fluorescence in the heart.

#### Tg(sox10:MRLC2-emGFP)

To generate the Sox10:MRLC2-emGFP construct (human MYL12B fused to Emerald GFP), the MRLC2 coding sequence was subcloned from the UAS-MLC2 plasmid, kindly provided by Adam Shellard (Roberto Mayor laboratory). The MRLC2 fragment was amplified by PCR and cloned downstream of the Sox10 promoter using the Sox10:mG backbone (kindly provided by Claudia Linker; Richardson et al., 2016) which contains Tol1 transposase sites and the sox10-4.9kb promoter region (Carney et al, 2006). Amplification and cloning were performed using the following primers: forward 5′-ACCTGTGGCCGCAGAACTAGTATGTCGAGCAAAAAGGCAAAGACC-3′ and reverse 5′-CAGTTCCTCTCCCTTTGAGACCATGTCATCTTTGTCTTTGGCTCCATG-3′.

To generate the MRLC2-emGFP fusion construct, the MRLC2 fragment was subsequently cloned in-frame with Emerald GFP using the 14xUAS-E1b:BGi cmlc2-eGFP backbone. Fusion to emGFP was achieved using the following primers: forward 5′-CATGGAGCCAAAGACAAAGATGACATGGTCTCAAAGGGAGAGGAACTG-3′ and reverse 5’-GGTGGCTGAGACTTAATTACTAGTTTACTTGTACAGCTCGTCCATGCC-3′. To generate the Sox10:MRCL2-emGFP transgenic line, 15 pg of pTol1-Sox10:MRCL-emGFP cmlc2-eGFP plasmid DNA was co-injected with 80 pg of Tol1 transposase mRNA into one-cell stage embryos. F1 founders were identified based on green fluorescence in the heart.

### Tg(sox10:EB3-GFP) line (sh691Tg)

We cloned the p5E -4725*sox10* zebrafish promoter (Dutton et al., 2008; Rodrigues et al., 2012), human EB3-EGFP (Stepanova et al., 2003) and p3E poly(A) sequences into pDestTol2pA3 using the Gateway Tol2 kit (Kawakami, 2007; Kwan et al., 2007). Embryos (AB strain) were injected and raised to adulthood. Transgenic progeny from one founder male crossed to AB were selected based on bright EB3-GFP expression and raised to adulthood to generate a stable line.

### Zebrafish maintenance

Zebrafish were maintained under standard, regulated conditions (Kimmel et al., 1995; Westerfield, 2007) at 28.5 °C on a 10 h light / 14 h dark cycle. All transgenic procedures and experimental work involving zebrafish were conducted in accordance with the Animal (Scientific Procedures) Act 1986, under Home Office Project Licence PP8606293 (PPL holder Elena Scarpa). Zebrafish work in Sheffield was performed under Home Office Project Licence PP2309825 (holder: Tanya T Whitfield).

Embryos were obtained from the following strains: wild type, TLF and AB strains; *Sox10:mG, Tg(–4.9sox10: Hsa.HIST1H2BJ-mCherry-2A-GLYPI-EGFP)* (Richardson et al., 2016); *Tg Sox10MRLC2-emGFP* (in this paper)*; Tg(sox10:EB3-GFP)* (this paper)*; Sox10:Kalt4, Tg(–4.9sox10: Hsa.HIST1H2BJ-mCherry-2A-Kalt4ER)* (Alhashem et al., 2021)*; Tg UAS-StableMark-Neongreen* (this paper) *; Tg UAS-EMTB-3xGFP* (this paper)*; Tg Sox10:mRFP* (fish kindly provided by Dr. Clara Mutschler)*; UAS:Dynamitin-CFP (*this paper). All embryos were raised in 1X E3 medium, prepared from a 60X stock solution containing 17.2 g NaCl, 0.76 g KCl, 2.9 g CaCl₂·H₂O, and 4.9 g MgSO₄. They were initially incubated at 28.5 °C until reaching 50% epiboly, after which the temperature was adjusted to either 22 °C (for the 14-somite stage) or 24 °C (for the 21-somite stage), depending on the desired developmental stage.

### In vivo imaging

Embryos were manually dechorionated using #3 Dumont forceps and mounted in 0.6% low-melting-point agarose prepared in E3 medium containing 0.002% MS-222 (Sigma, A5040-100G) on 35 mm glass-bottom dishes (Ibidi, 81158). Dishes were filled with E3 medium to maintain appropriate hydration and ensure embryo viability during long-term imaging (up to 8 hours).

For cranial neural crest imaging, embryos were mounted laterally with one side of the head positioned against the bottom of the dish. For mid-trunk imaging, embryos were also mounted laterally, with the trunk oriented against the bottom surface to maintain a flat imaging plane.

#### Neural crest cell migration imaging

Live imaging of neural crest cell migration (Sox10:mG and Sox10:Kalt4/UAS-Dynamitin-CFP) was performed using a PerkinElmer UltraVIEW spinning-disk inverted microscope equipped with an environmental chamber maintained at 28.5 °C. Imaging was carried out using a 30× UPlanSApo silicone immersion objective (Olympus), as previously described (Hakkinen et al., 2025). Z-stacks spanning 80–100 μm were acquired every 3 minutes over a total duration of 8 hours, using 20–30% laser power and 100 ms exposure time for both 488 nm and 561 nm channels.

#### Myosin and EB3-GFP imaging

Live imaging of MRLC2-emGFP and EB3-GFP was performed using a Nikon W1 SoRa spinning-disk inverted microscope equipped with an incubation chamber maintained at 28.5 °C and controlled using NIS-Elements software.

For myosin localisation (Sox10:MRLC2-emGFP) and centrosome localisation (EB3-GFP), imaging was performed using a 25×/1.05 NA CFI Plan Apo Lambda S silicone oil objective (Nikon; MRD73250). Z-stacks of ∼80 μm were acquired every 3 minutes over 8 hours using 2×2 binning and the sub-electron sensor system, with 5–10% laser power and single-frame exposure for both 488 nm and 561 nm channels.

For microtubule dynamics (*sox10*:EB3-GFP), imaging was performed using a 40×/1.25 NA CFI Plan Apo Lambda S silicone oil objective (Nikon; MRD73400). *Z*-stacks of ∼10 μm were acquired every 3 seconds for a total duration of 5 minutes, using 2×2 binning and the sub-electron sensor system, with 5–10% laser power and single-frame exposure for both 488 nm and 561 nm channels.

#### StableMark imaging

Live imaging of StableMark was performed using a Nikon Eclipse Ti microscope coupled to a Yokogawa CSU-X1 spinning-disk confocal system. Images were acquired with a 40× Nikon water immersion objective (DEMO) and a Photometrics Evolve camera. Z-stacks spanning 100 μm were acquired every 1-3 minutes over 6-8 hours, using 10–15% laser power for the 488 nm and 561 nm channels and an exposure time of 100 ms. Image acquisition was controlled using MetaMorph software.

### Laser ablation

#### Optical arrangement of confocal system

The confocal imaging module of the photomanipulation system is built based on a Nikon Ti2 inverted microscope coupled with an oa-the-shelf Confocal system (CrestOptics X-light V3). The setup is composed of a LDI-7 Laser Diode Illuminator (Oxxius Inc 89 North), a lens let array, a moveable spinning disk, two cameras (Photometrics Prime 95B), filters and lenses. The excitation light from the Laser illuminator is homogenised by the lens let array, passed through the excitation filter (Chroma ZET405/470/555/640x No.348855), focused on the spinning disk. After passing through the pin holes, the light is reflected by a dichroic mirror (Chroma ZT405/470/555/640rpc-UF2 No. 338528) and focused onto the microscope imaging plane. The emission light from a sample is collected by the same objective, follows a reciprocal path back through the spinning disk, passes through an emission filter (Chroma ZET405/470/555/640m-OD8 No. 34320), and is focused onto the camera. For dual-colour imaging, the emission light is split into GFP and mCherry channels via a dichroic mirror (Chroma T565lpxr-UF2 No. 347630) before reaching the cameras.

Individual bandpass filters (Chroma ET525/50m No. 345447 and ET605/70m No. 347621) are positioned before each camera to suppress stray light.

#### Photoablation system

The photoablation module is a custom developed sub-system composed of a femtosecond laser (Spectra-physics Mai Tai Deep sea Ti:Sapphire HP), a polarising beamsplitter, a Pockels cell (Lambda M350-105-BK-02), a beam expander, a spatial light modulator (Meadowlark 1920 x 1152), a set of filters and lenses. The ablation light is emitted from the Ti:Sapphire laser, polarised to plane waves, controlled by the Pockels cell to enable high-speed switching. Following beam expansion, the light is incident on a spatial light modulator for reshaping the beam profile. The modulated light is then relayed to the microscope’s image plane, allowing for precise spot or line ablation.

The laser wavelength was tuned to 980 nm to match the two-photon absorption spectrum of GFP. The laser output power was initially set to a low level (approximately 0.4 W, resulting in 0.13 W at the focal plane) to minimise direct thermal damage to cells. Once the ablation site on the microtubule bundle is precisely identified, the shutter was open for 100 ms to allow the ablation light to scan along a line (5 µm in length) on the specimen. If no obvious ablation occurs, the laser power was increased incrementally in steps of 0.05 W, up to a maximum of 1.0 W, until the microtubule bundle is successfully ablated. The entire ablation process is controlled by a custom-developed LabVIEW programme, while the confocal system is controlled by MetaMorph (Molecular Device) to continuously image the ablation and monitor the changes in nuclear shape.

### Drug treatments

#### Pharmacological inhibitions

Embryos were collected and maintained in E3 medium until the 18-somite stage (at 24°C for trunk region) or 10-somite stage (at 22°C for cranial region). At this stage, pharmacological treatments were applied to inhibit contractility or dynein function, and embryos were maintained in drug-containing medium throughout the duration of imaging.

To inhibit ROCK, embryos were treated with 10, 20, or 50 μM of the ROCK inhibitor Y-27632 (Cat# HY-10071, MedChemExpress). The inhibitor was prepared as a 5 mg/mL stock solution in DMSO (D2650, Sigma) and diluted in E3 medium prior to use. For all experiments presented, a concentration of 50 μM was used, as this was the highest dose required to fully disrupt contractility and membrane retraction. Control embryos were treated with 1% DMSO under identical conditions at 28.5 °C. Time-lapse imaging was performed for 8 hours with the drug present throughout acquisition.

To inhibit dynein function, embryos were treated with 15, 25, 30, or 50 μM Ciliobrevin D (Cat# HY-122632, MedChemExpress). A concentration of 15 μM was selected for subsequent experiments, as it represented the lowest dose that produced a consistent and detectable phenotype. Control embryos were treated with 1% DMSO (Sigma-Aldrich) under the same conditions.

#### Gene expression induction

Sox10:Kalt4 embryos were outcrossed with UAS responder lines (UAS:StableMark, UAS:EMTB-3xGFP, and UAS:Dynamitin-CFP). Gene expression was induced by incubating embryos in tamoxifen (Sigma-Aldrich, H7904) added to E3 medium at the shield stage. Final concentrations of 0.5 μM were used for StableMark expression, and 0.25 μM for EMTB-3xGFP and Dynamitin-CFP expression.

Following induction, embryos were maintained in tamoxifen-containing medium at stage-specific temperatures (22 °C for cranial imaging and 24 °C for trunk imaging) to allow proper development and transgene expression. Prior to mounting, embryos were washed to remove tamoxifen and embedded in agarose prepared in E3 medium without tamoxifen.

Time-lapse imaging was then performed at 28.5 °C for a total duration of 8 hours.

### Immunofluorescence

Immunostaining was performed using a modified zebrafish deyolking protocol (Frommelt et al., 2023) to improve antibody penetration for nuclear protein detection. Embryos at the 22-somite stage (trunk) and 14-somite stage (cranial) were manually dechorionated and fixed in 1% paraformaldehyde (PFA) in 1× PBS for 2 hours at room temperature with gentle agitation.

Following fixation, embryos were washed three times in 1× PBS containing Triton X-100 (0.1% for trunk; 0.01% for cranial samples; T6767, Sigma). The yolk was then carefully removed using a fine needle and forceps in 1× PBS. Embryos were subsequently post-fixed in 4% PFA in 1× PBS for an additional 2 hours at room temperature with gentle agitation.

Permeabilization was carried out with three washes in PBS containing Triton X-100 (0.5% for trunk; 0.1% for cranial) at room temperature and gentle agitation. Embryos were then blocked for 2 hours at room temperature in blocking solution (trunk: 0.5% Triton X-100 in PBS, 4% normal goat serum, 2% DMSO; cranial: 0.1% Triton X-100 in PBS, 4% normal goat serum, 2% DMSO).

Primary antibodies were diluted in blocking solution and incubated with the embryos for 48 hours at 4 °C in gentle agitation. Samples were then washed 10 times for 5 minutes each in PBS 1X containing Triton X-100 (0.5% for trunk; 0.1% for cranial). Secondary antibodies were applied in blocking solution for 2 hours at room temperature or overnight at 4 °C.

After secondary incubation, embryos were washed 10 times for 5 minutes each in PBS with Triton X-100 (0.5% for trunk; 0.1% for cranial), mounted on glass slides in 75% glycerol/PBS, borders were sealed and the slides were imaged using an Olympus FV3000 confocal microscope with a 30× silicone immersion objective.

The following primary antibodies were used: anti-γ-tubulin (mouse monoclonal, T6557, MilliporeSigma, 1:100); anti-DYNC1H1 (rabbit polyclonal, 12345-1-AP, Proteintech, 1:100); anti-RFP (rat monoclonal, 5F8, ChromoTek, 1:200); anti-mCherry (rabbit polyclonal, GTX128508, GeneTex, 1:200); anti-mCherry (chicken polyclonal, AB205402, Abcam, 1:200); anti-lamin B2 (mouse monoclonal, AB8983, Abcam, 1:200); and anti-GFP (chicken polyclonal, AB13970, Abcam, 1:200).

Secondary antibodies were used at a 1:500 dilution: Alexa Fluor 488 goat anti-chicken (A11039, Invitrogen); Alexa Fluor 488 chicken anti-mouse (A21200, Invitrogen); Alexa Fluor 546 goat anti-rat (A11081, Invitrogen); Alexa Fluor 546 goat anti-chicken (A11040, Invitrogen); Alexa Fluor 546 goat anti-rabbit (A11035, Invitrogen) and Alexa Fluor 647 Fab goat anti-rabbit (A48285, Invitrogen).

### Image Analysis

#### 1D Kinematics Analysis of nuclear and membrane motion

**Methods Figure 1.**
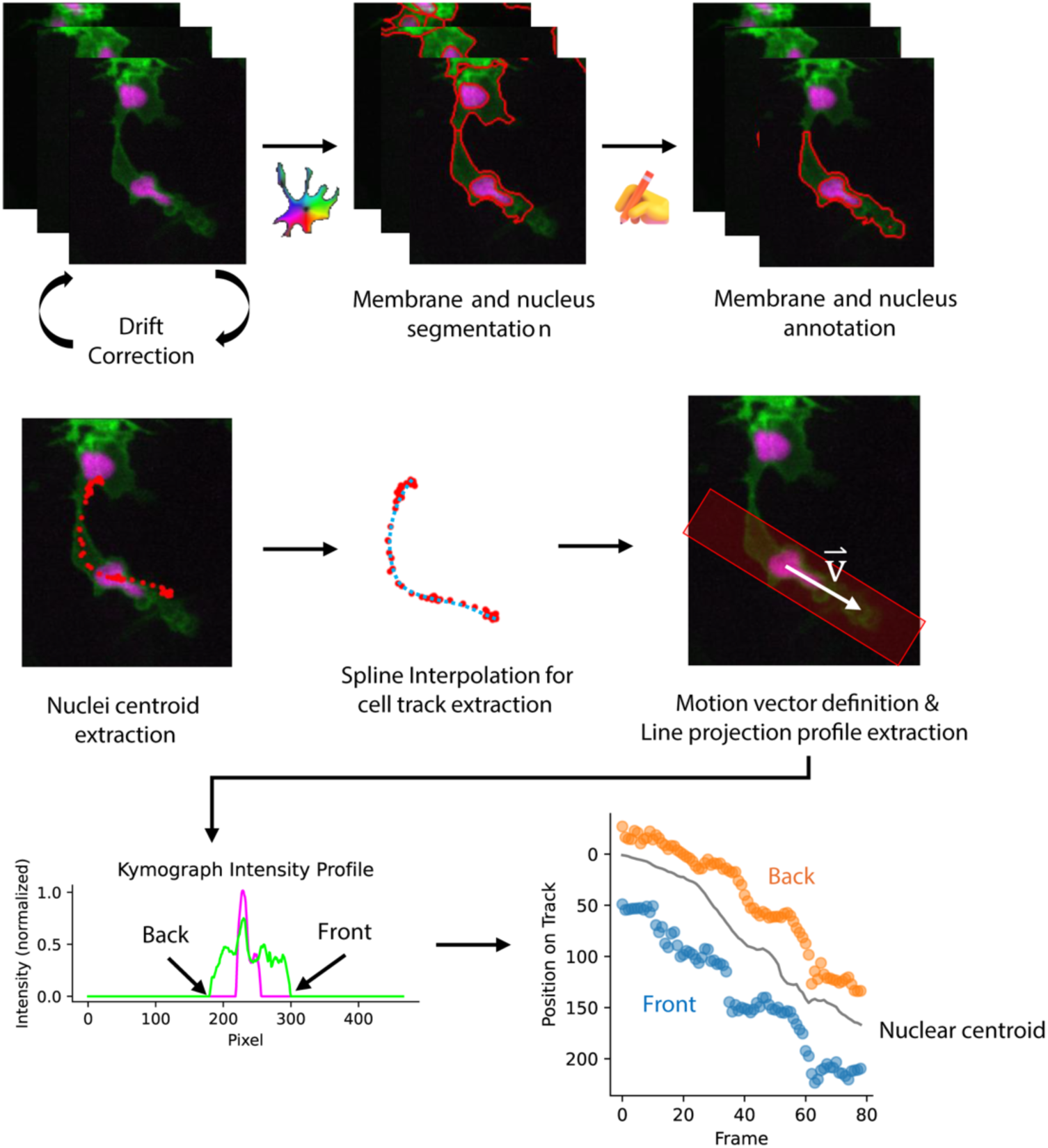
Image analysis pipeline for kymograph generation and quantification of nuclear and membrane dynamics. Time-lapse image sequences were first corrected for stage drift before membrane and nuclear structures were segmented and manually curated. Nuclear centroids were extracted from the segmented nuclei and linked across frames using spline interpolation to generate a continuous cell trajectory. A motion vector (v) was defined along the migration path, and line projection profiles were extracted along the cell axis. Intensity profiles from each projection were used to identify the front and back membrane positions relative to the nuclear centroid. These measurements were compiled over time to generate kymographs and quantify the dynamics of the cell front, back, and nucleus during migration.

#### Extraction of cell migration features

To quantify the dynamic morphological and migratory behaviour of neural crest cells, we developed a multi-step computational pipeline implemented in Python. Initial image stacks were segmented using *Cellpose*3 to generate separate masks for the nuclei and cell membranes. Segmented masks were manually inspected, refined and tracked using a custom *Napari*-based interface to ensure temporal consistency and correct any segmentation artefacts.

#### Image registration and drift correction

To account for global sample drift during long-term imaging, we applied a phase-cross correlation method. The nuclear channel stack was used as a reference stack to compute the cumulative translation between successive frames using Fast Fourier Transforms (FFT). These calculated shifts were then applied to both the membrane and nuclei image stacks and their corresponding segmentation masks to ensure spatial alignment across the duration of the experiment.

#### Morphological and kinematic analysis

For each tracked cell, we extracted primary morphological features of the membrane and nucleus over time, including area (*A*), perimeter (*P*), and centroid coordinates (*x, Y*), using the *regionprops_table* function from the scikit-image library. Circularity was calculated as 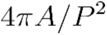. Instantaneous migration speed was defined as the Euclidean distance between centroids in adjacent frames: 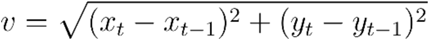 where *<u>t</u>* represents the frame index.

#### Trajectory reconstruction and 1D projection

To minimise the impact of nuclear “jittering” caused by both biological fluctuation and stochastic observation noise, we reconstructed the principal migration path for each cell using a B-spline interpolation. A smoothed spline was fit to the raw nuclear centroid 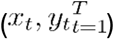 using *scipy.interpolate.make_splprep* with a smoothing condition (*s* = 100).

The raw centroid at each time point was then projected onto this 1-pixel spaced spline using a k-dimensional tree (cKDTree) search to determine its position on the track. To further refine these coordinates, a temporal rolling average (window=5 frames) was applied to the projected indices.

#### Directional line profiling and boundary extraction

To analyse membrane dynamics relative to nuclear motion, we extracted 1D intensity profiles along the cell’s current direction of travel. The motion vector 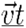 at time *<u>t</u>* was defined as: 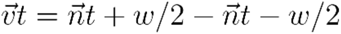 where 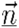 is the nuclear centroid and *<u>w</u>* is the window size (set to 16 frames) to capture the persistent direction of motion.

Centring on the nucleus, we extracted a line profile of 50 pixels in width along this vector. The membrane boundaries (membrane front and back) were identified as the first and last points along this profile where the membrane segmentation mask signal was present. These boundary locations were then mapped back to the 1D spline coordinate system, allowing for the construction of kymographs that represent the relative expansion and contraction of the cell body along its migration axis. All extracted parameters were exported for downstream statistical analysis.

#### Microtubules Dynamics and Comets orientation

The *in vivo* movies for EB3-GFP dynamics were analysed automatically through MATLAB using the plusTipTracker software (Applegate et al., 2011), as previously described (Villaseca et al., 2025). The following parameters were applied: 8 frames as the maximum gap length, 3 frames as the minimum track length, 5-10 pixels as the search radius range, 30 as the maximum forward angle, 10 as the maximum backward angle, 1.5 as the maximum shrinkage factor and a 1-pixel fluctuation radius. Frame rate was 3 seconds and pixel size was 0.116 nm. Comet lifetime, growth speed and growth length were analysed for cranial and trunk neural crest cells. Comet orientation was also obtained from plusTipTracker quantification, and the graphs were made in R using an adapted code for R-Studio (adapted from Scarpa et al., 2018).

#### Stable microtubules distribution

To quantify the *in vivo* distribution of StableMark signal, the Plot Profile function in FIJI (NIH, https://imagej.net/fiji) was used to analyse stable microtubule localisation at the point of maximum nuclear deformation (defined as the largest decrease in circularity), as previously described (Hakkinen et al., 2025).

Time-lapse movies were first corrected for XYZ drift using the Registration plugin in FIJI, followed by generation of maximum intensity projections along the Z-axis. Nuclear segmentation was performed using StarDist, and the time point of maximum deformation was identified manually based on the minimum circularity value.

For each time point within a deformation event, a 10-pixel-wide line was drawn from the rear to the front of the nucleus along the direction of migration. The Plot Profile function was then used to measure the distribution of StableMark fluorescence intensity along this line. Nuclear signal distribution was analysed in parallel for comparison.

Fluorescence intensity values were normalised to the average intensity of each individual line profile. To account for variations in nuclear length over time, profiles were rescaled by binning the data into equal segments, where 0% corresponds to the rear and 100% to the front of the nucleus relative to the direction of migration.

#### Centrosome indentation and orientation

To quantify centrosome localisation, we analysed two parameters: centrosome indentation and centrosome orientation, both associated with the direction of cell migration and nuclear deformation.

Using FIJI, a 1-pixel-wide line was drawn from the nuclear centroid (obtained from segmentation) to the centrosome position to measure the centrosome–centroid distance (d). The nuclear radius (r) was defined as the distance from the nuclear centroid to the nuclear envelope. A ratio (d/r) was then calculated for each measurement.

A ratio below 1 indicates that the centrosome is located within the nuclear radius, consistent with positive nuclear indentation, whereas a ratio greater than 1 indicates that the centrosome lies outside the nuclear boundary.

To validate centrosome indentation, the d/r ratio was correlated with the point of maximum nuclear deformation, defined as the time point exhibiting the lowest nuclear circularity.

For centrosome orientation, the angle between the centrosome position and the nuclear axis aligned with the direction of migration was measured. Angles were determined before, during, and after nuclear deformation. The centrosome angle was subtracted from the nuclear angle, and the absolute value of this difference was used to define centrosome orientation relative to the direction of migration.

Centrosome orientation was categorized as follows: angles between 0° and 15° indicate orientation toward the front of migration; angles between 15° and 45° indicate lateral orientation; and angles greater than 45° indicate lateral-to-rear orientation relative to the migration direction.

#### Nucleus recoil

Laser ablation experiments were imaged at 1-second intervals for up to 3 minutes. Image sequences were analysed using the Dynamic Reslice function in FIJI to visualise recoil dynamics over time.

Nuclear recoil was quantified following segmentation with the StarDist plugin. Changes in nuclear circularity were measured over time in trunk neural crest (tNC; confined) and cranial neural crest (cNC; non-confined) cells. Nuclear circularity was obtained every second until the end of the movie after microtubules ablation.

For tNC cells, nuclei were further classified as deformed or non-deformed based on their initial morphology prior to ablation.

#### Dynein and LaminB2 localisation

Cytoplasmic dynein heavy chain and Lamin B2 localisation were analysed by immunostaining in Sox10:KalTA4 zebrafish embryos expressing H2B-mCherry under the Sox10 promoter.

Confocal Z-stacks were reorganised using the Reorder Hyperstack function in FIJI. Nuclear segmentation was performed using H2B-mCherry signal and the StarDist plugin. Segmented nuclei were thresholded and eroded three times to generate reduced nuclear masks (“smaller nuclei”), which were added to the ROI Manager and annotated according to cell type (non-confined cranial neural crest cells, non-confined trunk neural crest pre-migratory cells, and confined trunk neural crest migratory cells).

To analyse perinuclear localisation, reduced nuclear ROIs were expanded into 7-pixel-wide outlines corresponding to the nuclear envelope and added to the ROI Manager. These ROIs were then applied to the dynein heavy chain and Lamin B2 channels to quantify fluorescence intensity at the nuclear periphery.

For cytoplasmic measurements, original (non-eroded) nuclear ROIs were used to generate 7-pixel-wide outlines representing the cytoplasmic region surrounding the nucleus. These ROIs were similarly annotated and applied to the corresponding fluorescence channels to quantify cytoplasmic signal intensity.

For paired analyses, fluorescence intensity values were normalised by the area of each ROI (perinuclear and cytoplasmic). For comparative analyses, a perinuclear-to-cytoplasmic intensity ratio was calculated. A ratio greater than 1 indicates enrichment at the nuclear envelope, whereas a ratio equal to or below 1 indicates equal or higher signal in the cytoplasm.

### Statistical data analysis

The statistics used in this paper, *p*-values, number of data points, number of cells, number of embryos and number of experiments are all mentioned in the figure legends. Statistical tests were performed using GraphPad Prism version 10.4.1 (Windows). Data plots were generated in R using R Studio scripts, Python and GraphPad Prism.

## Author contributions

Conceptualisation and experimental design: SV, ES. Experimental work: SV, MP, CZ, ZA, HMH. Segmentation and quantitative kinematics pipeline: CYH. Generation of *Tg(sox10:EB3-GFP)* line: SB, MM, TTW. Development and implementation of the photoablation microscope system; CZ, ML .Generation of Tg(sox10:MRCL2-emGFP) line: MDLB. Generation of Tg(UAS-Dynamitin-CFP) line: AH. Generation of Tg(UAS-StableMark); Tg(UAS-EMTB-3x-GFP) lines: DEZ, SV. Manuscript writing: SV, ES. Review of manuscript: SB, TTW, MP, CYH. Funding: ES, TTW.

## Acknowledgements

Scarpa lab: ES is the recipient of a Royal Society Dorothy Hodgkin Fellowship (DHF\R1\201118, (RF\ERE\210154, RF\ERE\231096); MRC New Investigator Grant to ES (MR/W024519/1). Isaac Newton Trust Grant (2507.07(e) to ZA and ES. MRC Mid-Range Equipment grant to ES (MC_PC_ APP23060). Whitfield lab: BBSRC Project Grant (BB/S007008/1) to TTW and SB, and a Wellcome Trust Investigator Award (224355/Z/21/Z) to TTW. Support through Isaac Newton Trust Grant (25.07(c)) for CZ. We thank Jon Howe, Ben Sutcliffe and Antonina Kruppa of the Light Microscopy Facility of the Microscopy Bioscience Platform for technical support and insightful advice. We thank Raymond Wightman of the Sainsbury Laboratory Microscopy Facility for kindly providing access to the Nikon Eclipse Ti microscope. We also thank Fiona Morgan and Ewa Paluch for their technical support with the PerkinElmer UltraVIEW spinning-disk microscope. We thank Claudia Linker for sharing the *sox10:mG, sox10-Kalt4 and Sox10:H2B-Dendra* zebrafish lines and her expertise, time, reagents and data. We thank Gökçe Agsu for insightful discussions. We thank Clara Mutschler and Peter Arthur-Farraj for sharing the sox10:mRFP transgenic. We thank Reinhard Köster for the gift of UAS-Dynamitin-CFP. The Tol2Kit was a gift from Koichi Kawakami. We thank Lukas Kapitein for gifting the StableMark plasmid (Addgene #174649), Willliam Bement for the gift of the pCS2+-EMTB-3X-EGFP plasmid (Addgene #26741), Adam Shellard for the gift of UAS-MRLC2-GFP. We are grateful to Caren Norden and Ben Steventon for critical reading of the manuscript.

---

**Figure S1.**
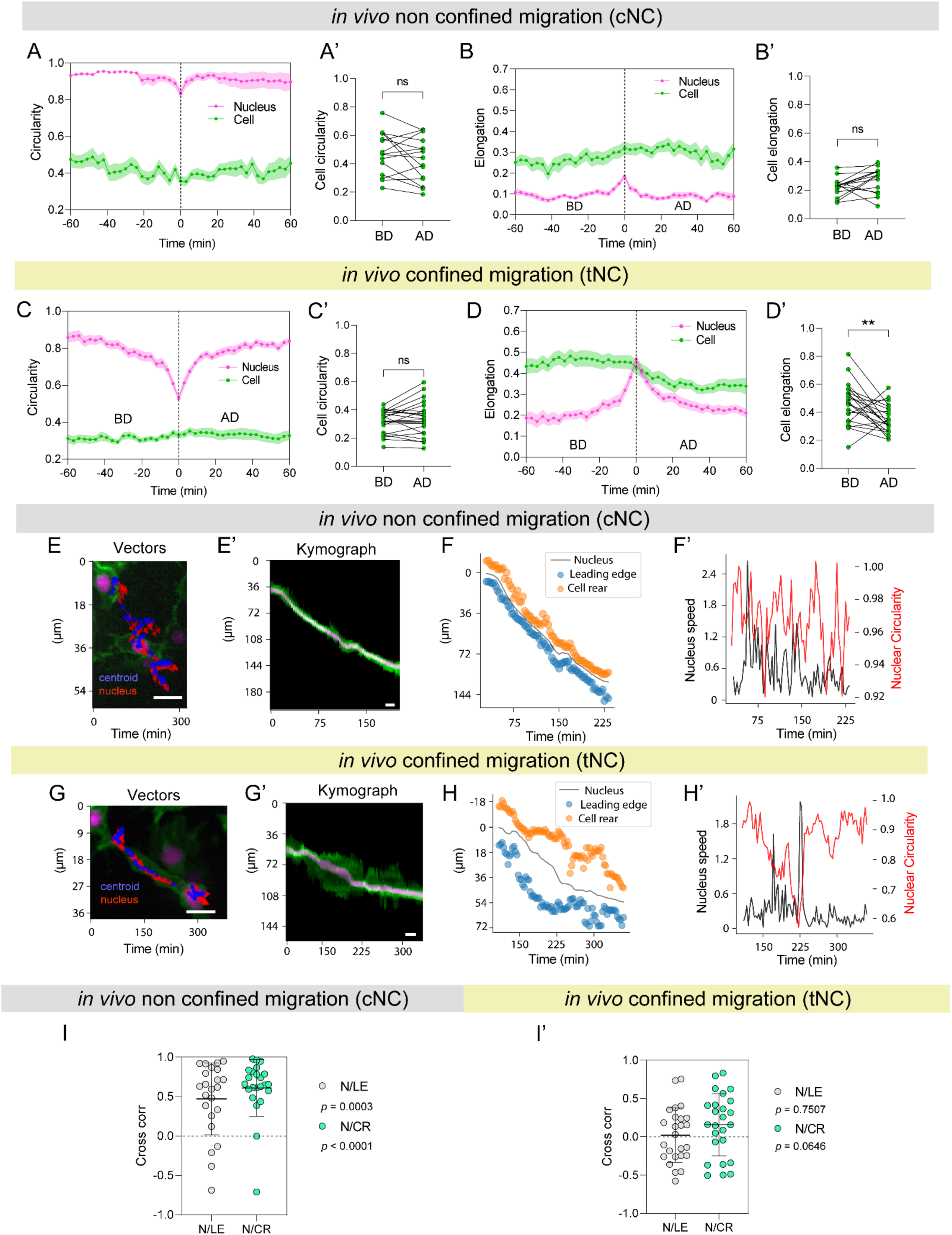
Nuclear and cell morphodynamics reveal uncoupled nucleokinesis during confined trunk neural crest migration *in vivo*. (A) Mean nuclear (magenta) and cell (green) circularity over time in non-confined cranial neural crest (cNC) cells, aligned to the point of maximum nuclear deformation (time = 0). Data span 60 min before and after alignment. BD, before deformation; AD, after deformation. *n* = 23 cells from 4 embryos across 3 experiments. Error bars represent SEM. (A’) Comparison of cell circularity before (BD) and after (AD) nuclear deformation in non-confined cNC cells using a two-tailed paired *t*-test. *n* = 15 cells from 4 embryos across 3 experiments; *p* = 0.1672. For paired comparisons, only cells with matched measurements before and after nuclear deformation were included. (B) Mean nuclear (magenta) and cell (green) elongation over time in non-confined cNC cells, aligned to the point of maximum nuclear deformation (time = 0). Data span 60 min before and after alignment. BD, before deformation; AD, after deformation. *n* = 23 cells from 4 embryos across 3 experiments. Error bars represent SEM. (B’) Comparison of cell elongation before (BD) and after (AD) nuclear deformation in non-confined cNC cells using a two-tailed paired *t*-test. *n* = 15 cells from 4 embryos across 3 experiments; *p* = 0.0553. For paired comparisons, only cells with matched measurements before and after nuclear deformation were included. (C) Mean nuclear (magenta) and cell (green) circularity over time in confined trunk neural crest (tNC) cells, aligned to the point of maximum nuclear deformation (time = 0), defined as the time point of minimum nuclear circularity. Data span 60 min before and after alignment. BD, before deformation; AD, after deformation. *n* = 25 cells from 6 embryos across 5 experiments. Error bars represent SEM. (C’) Comparison of cell circularity before (BD) and after (AD) nuclear deformation in confined tNC cells using a two-tailed paired *t*-test. *n* = 23 cells (valid BD and AD times) from 6 embryos across 5 experiments; *p* = 0.2819. For paired comparisons, only cells with matched measurements before and after nuclear deformation were included. (D) Mean nuclear (magenta) and cell (green) elongation over time in confined tNC cells, aligned to the point of maximum nuclear deformation (time = 0). Data span 60 min before and after alignment. BD, before deformation; AD, after deformation. *n* = 25 cells from 6 embryos across 5 experiments. Error bars represent SEM. (D’) Comparison of cell elongation before (BD) and after (AD) nuclear deformation in confined tNC cells using a two-tailed Wilcoxon test. *n* = 46 cells from 4 embryos across 3 experiments; *p* = 0.0030. For paired comparisons, only cells with matched measurements before and after nuclear deformation were included. (E–F′) Representative quantification workflow for a single non-confined cNC cell migrating in vivo. (E) Schematic summarizing migration and the analysis pipeline. Scale bar, 10 µm. (E′) Kymograph of membrane (green) and nuclear (magenta) intensity profiles aligned to the nuclear centroid. Scale bar, 10 µm. (F) Temporal trajectories of nuclear position (black), leading edge (blue), and cell rear (orange) throughout migration. (F′) Representative analysis of nuclear circularity (red) and nuclear speed (black), showing no temporal association between nuclear deformation and nuclear movement. (G–H′) Representative quantification workflow for a single confined tNC cell migrating in vivo. (G) Schematic summarizing migration and the analysis pipeline. Cell rear, nuclear centroid, and leading edge positions were tracked over time. Red arrows indicate nuclear movement and blue dots indicate the cell centroid. Scale bar, 10 µm. (G′) Kymograph of membrane (green) and nuclear (magenta) intensity profiles aligned to the nuclear centroid along the migration axis. Scale bar, 10 µm. (H) Temporal trajectories of nuclear position (black), leading edge (blue), and cell rear (orange) throughout migration. (H′) Representative analysis showing nuclear circularity (red) and nuclear speed (black) over time, illustrating that the point of minimum nuclear circularity coincides with maximal nuclear speed. (I) Cross-correlation analysis between nucleus (N) and leading edge (LE), and between nucleus (N) and cell rear (CR), in non-confined cNC cells. Correlation coefficients were tested against a theoretical median of 0 using a one-sample Wilcoxon test. Both N–LE (*p* = 0.0003) and N–CR (*p* < 0.0001) differed significantly from zero, indicating coordinated nuclear and cell edge dynamics. *n* = 23 cells from 4 embryos across 3 experiments. (I’) Cross-correlation analysis between nucleus (N) and leading edge (LE), and between nucleus (N) and cell rear (CR), in confined tNC cells. Correlation coefficients were tested against a theoretical value of 0 (no correlation) using a one-sample *t*-test. Values approaching 1 indicate positive correlation and values approaching 0 indicate lack of correlation. Neither N–LE (*p* = 0.7507) nor N–CR (*p* = 0.0646) differed significantly from zero, indicating uncoupled nuclear and cell edge dynamics. *n* = 25 cells from 6 embryos across 5 experiments.

**Figure S2.**
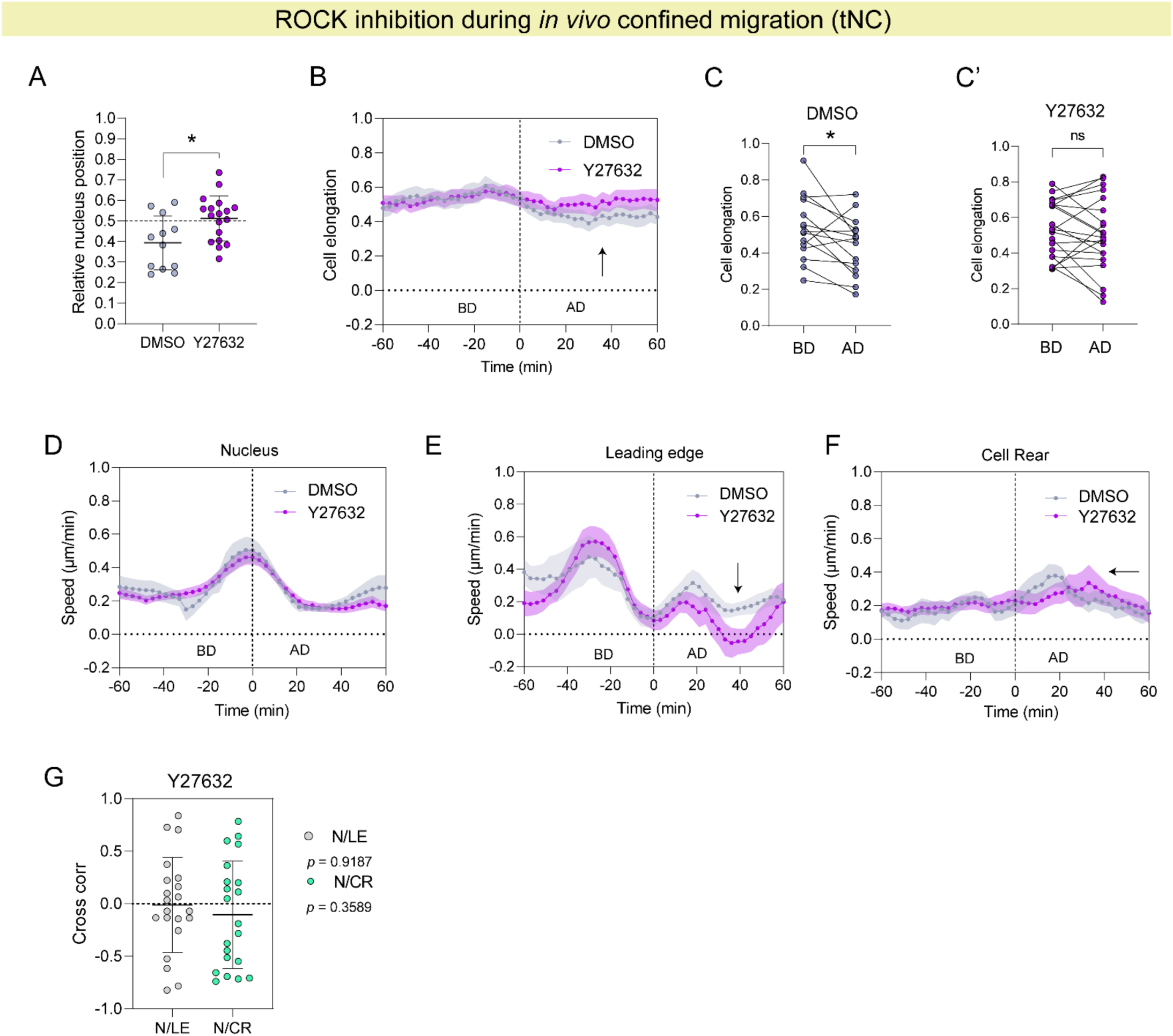
Nuclear and cell morphodynamics under ROCK inhibition reveal persistent uncoupling of nuclear and cell edge dynamics. (A) Comparison of relative nuclear position in trunk neural crest (tNC) cells between DMSO- and Y27632-treated embryos during the period 40–60 min after maximum nuclear deformation. Under ROCK inhibition, nuclei remained positioned closer to the cell centre following the nuclear translocation period compared with DMSO controls. Two-tailed unpaired *t*-test. DMSO: *n* = 15 cells from 4 embryos across 3 experiments. Y27632: *n* = 21 cells from 8 embryos across 4 experiments. *p* = 0.0117. (B) Mean cell elongation over time in DMSO- and Y27632-treated embryos, aligned to the point of maximum nuclear deformation (time = 0). Data span before (BD) and after deformation (AD). The black arrow highlights the interval during which Y27632-treated cells remain elongated (after maximum nuclear deformation) compared with DMSO controls. DMSO: *n* = 15 cells from 4 embryos across 3 experiments. Y27632: *n* = 21 cells from 8 embryos across 4 experiments. Error bars represent SEM. (C, C’) Two-tailed paired *t*-test comparing cell elongation before deformation (BD) and after deformation (AD) in DMSO (C) and Y27632 (C′) conditions. DMSO showed a significant change in elongation (*p* = 0.0284), whereas no significant difference was detected following ROCK inhibition (*p* = 0.4105). (D-F) Mean temporal dynamics of nuclear speed (D), leading-edge speed (E), and cell rear speed (F) in DMSO-and Y27632-treated cells aligned to maximum nuclear deformation (time = 0). BD, before deformation; AD, after deformation. Black arrows indicate delayed coordination of leading-edge dynamics after nuclear deformation (E) and delayed cell rear dynamics (F) under Y27632 treatment. (G) Cross-correlation analysis between nucleus (N) and leading edge (LE), and between nucleus (N) and cell rear (CR), in confined tNC cells treated with Y27632. Correlation coefficients were tested against a theoretical value of 0 (no correlation) using a one-sample *t*-test. Values approaching 1 indicate positive correlation, whereas values approaching 0 indicate lack of correlation. Neither N–LE (*p* = 0.9187) nor N–CR (*p* = 0.3589) differed significantly from zero, indicating that ROCK inhibition does not restore coupling between nuclear movement and cell edge dynamics. *n* = 21 cells from 8 embryos across 4 experiments.

**Fig. S3.**
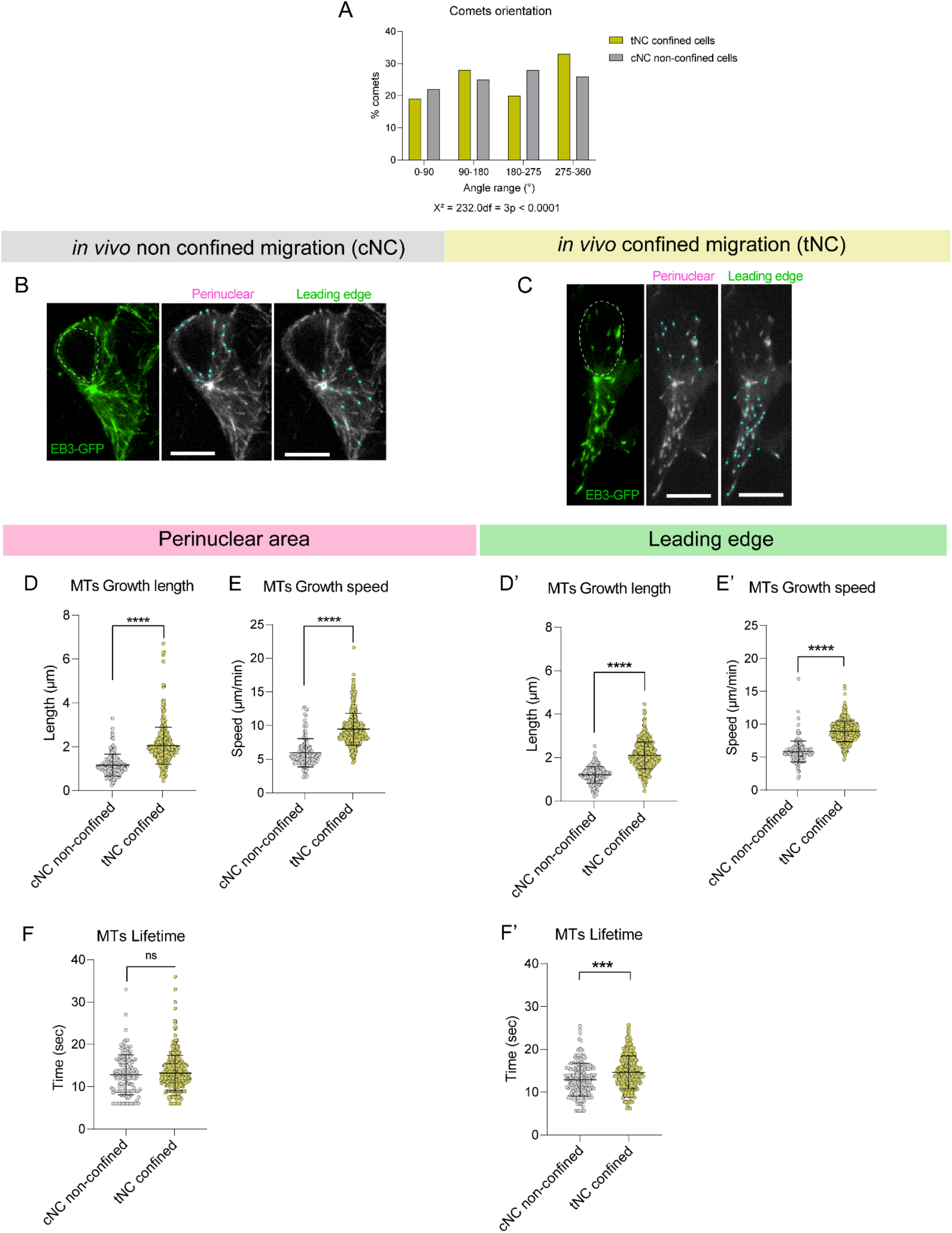
Confined migration alters subcellular microtubule growth dynamics *in vivo*. (A) Distribution of comet orientation angles in non-confined cNC cells (grey bars) and confined tNC cells (yellow bars). Orientation angles obtained from comet analysis were converted into a 0–360° angular representation and grouped into four angular sectors (0–90°, 90–180°, 180–275°, and 275–360°). Bars represent the percentage of total valid orientation measurements within each angular interval. A total of *n* = 6,804 valid comet orientations for cNC cells and *n* = 10,518 valid comet orientations were analysed for tNC cells. cNC cells displayed a more evenly distributed orientation profile across angular sectors, with a modest enrichment in the 180–275° sector (28.2%), followed by 275–360° (25.5%), 90–180° (24.6%), and 0–90° (21.7%). cNC: n = 27 cells, 6 embryos, 2 experiments. In contrast, in tNC the distribution showed a preferential accumulation of orientations within the 275–360° sector (33.5%), followed by 90–180° (27.6%), 180–275° (20.2%), and 0–90° (18.6%). n = 21 cells tNC cells, 6 embryos, 3 experiments. Angular sector occupancy was compared between confined tNC and non-confined cNC cells using a χ² test of independence. The distribution of comet orientations differed significantly between conditions (χ² = 232.0, df = 3, *p* < 0.0001), with confined tNC cells displaying enrichment in the 275–360° sector. (B, C) Representative images of EB3-GFP positive cNC cells (C) and tNC cells (D) where a ROI was defined at the perinuclear area and the leading edge. The comets were automatically detected by the PlusTipTracker software as cyan dots and followed for 5 minutes, 3 seconds each frame. (D-F) EB3-GFP comet growth length (D), speed (E) and lifetime (F) in cNC non-confined cells (grey) versus tNC confined cells (yellow) at the Perinuclear area only. Non-confined cNC: n = 27 cells, 1513 perinuclear comets, 6 embryos, 2 experiments. Confined tNC: n = 21 cells, 3825 perinuclear comets, 6 embryos, 3 experiments. Two tailed Mann Whitney test. **** p < 0.0001; ns p = 0.3172. Each dot represents one comet. (D’-F’) EB3-GFP comet growth length (D), speed (E) and lifetime (F) in non-confined cNC cells (grey) versus confined tNC cells (yellow) at the Leading edge only. Non-confined cNC: n = 27 cells, 2656 leading edge comets, 6 embryos, 2 experiments. Confined tNC: n = 21 cells, 4427 leading edge comets, 6 embryos, 3 experiments. Two tailed Mann Whitney test. **** p < 0.0001; *** p = 0.0009. Each dot represents one comet.

**Figure S4.**
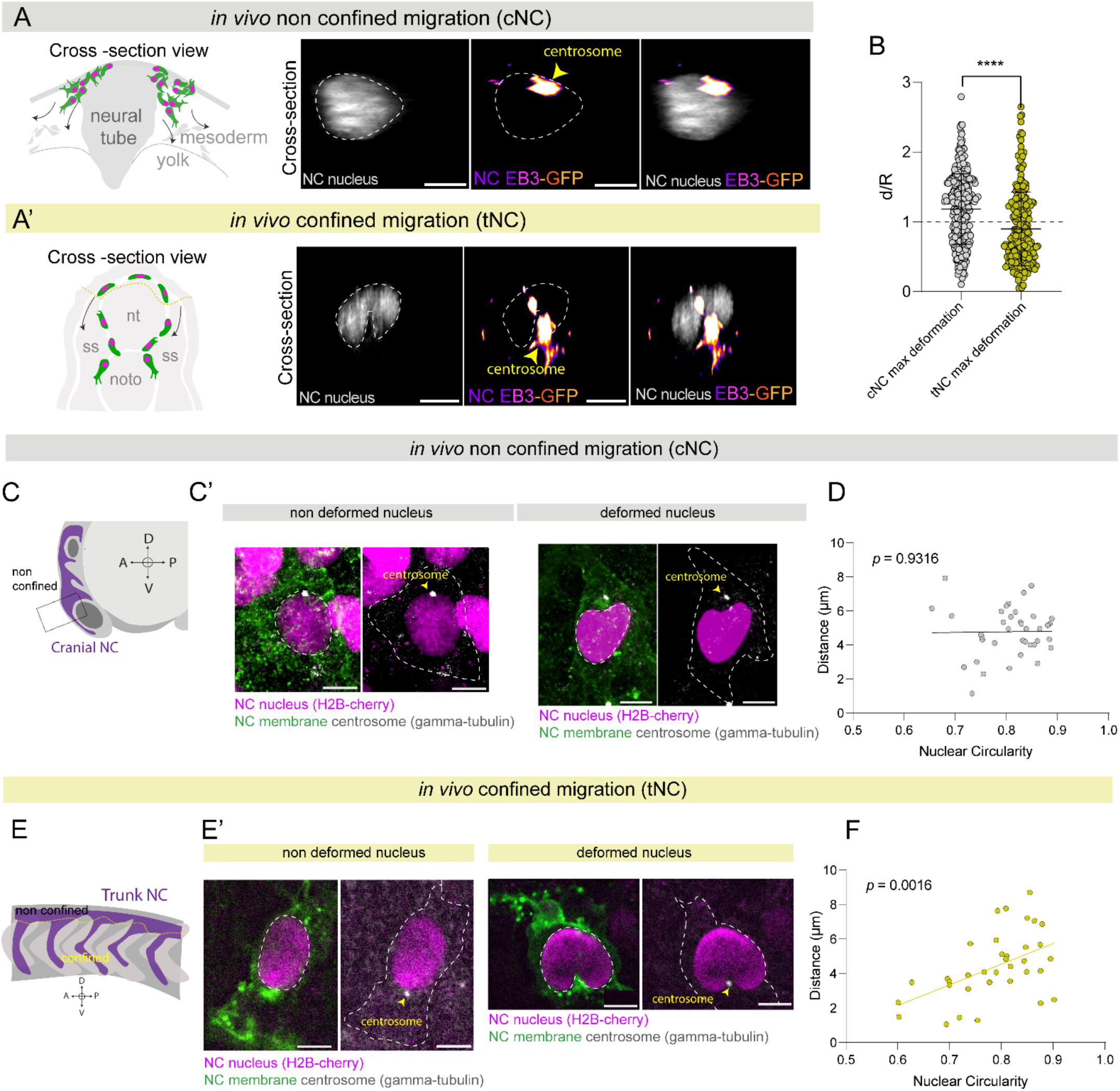
Centrosome–nucleus coupling correlates with nuclear deformation in confined but not non-confined migration *in vivo*. (A) Cross-section view showing that at the first panel a schematic of cranial neural crest (cNC) cells migrating within a non-confined environment. Second panel shows cross section view for a representative cNC cell expressing EB3GFP (in pseudocolor) and H2B-Cherry (in grey) with no nuclear indentation evidence nor nuclear deformation in the Z plane. (A’) Cross-section view showing that at the first panel a schematic representation of the mid-tNC illustrating the space available between the tissue for the transition from non-confined (premigratory) to confined (migratory) tNC cells. Second panel showing a cross section view for a representative tNC cell expressing EB3-GFP (in pseudocolor) and H2B-Cherry (in grey) which evidence positive nuclear indentation of the centrosome and deformation of the nucleus in the Z plane. (B) Quantification of nuclear indentation at the point of maximal deformation (defined by a sharp decrease in nuclear circularity) compared between tNC and cNC cells. tNC: *n* = 17 cells, 9 embryos, 3 experiments, 240 time points. cNC: *n* = 21 cells, 7 embryos, 3 experiments, 300 time points; Two-tailed Mann–Whitney test; ****, *p* < 0.0001. Each dot represents one time point. (C) Schematic of cranial neural crest (cNC) cells in a 14-somite-stage zebrafish embryo, migrating within a non-confined environment. (C’) Representative immunofluorescence images of polarized (E) and deformed (E’) cNC cells. Nuclei are shown in magenta and centrosomes (γ-tubulin) in grey. Yellow arrowheads indicate the position of the centrosome within the cell. Scale bar, 5 µm. (D) Linear regression analysis in cNC cells showing no correlation between centrosome distance and nuclear circularity. *n* = 38 cells; *p* = 0.9316, not significantly different from zero. (E) Schematic of the mid-tNC (between the 7th and 10th somites), illustrating the transition from non-confined (premigratory) to confined (migratory) environments. (E’) Representative immunofluorescence images of non-deformed and deformed nucleus of tNC cells from Sox10:mG transgenic embryos. Nuclei are shown in magenta, membrane is shown in green and centrosomes (γ-tubulin staining) in grey. Yellow arrowheads indicate the position of the centrosome within the cell. Scale bar, 5 µm. (F) Linear regression analysis of centrosome distance from the nuclear centroid versus nuclear circularity in fixed tNC cells. Higher nuclear circularity (≈1) correlates with increased centrosome distance, whereas decreased circularity (greater deformation) correlates with centrosome proximity to the nuclear centroid. *n* = 36 cells; *p* = 0.0016, significant deviation from zero.

**Figure S5.**
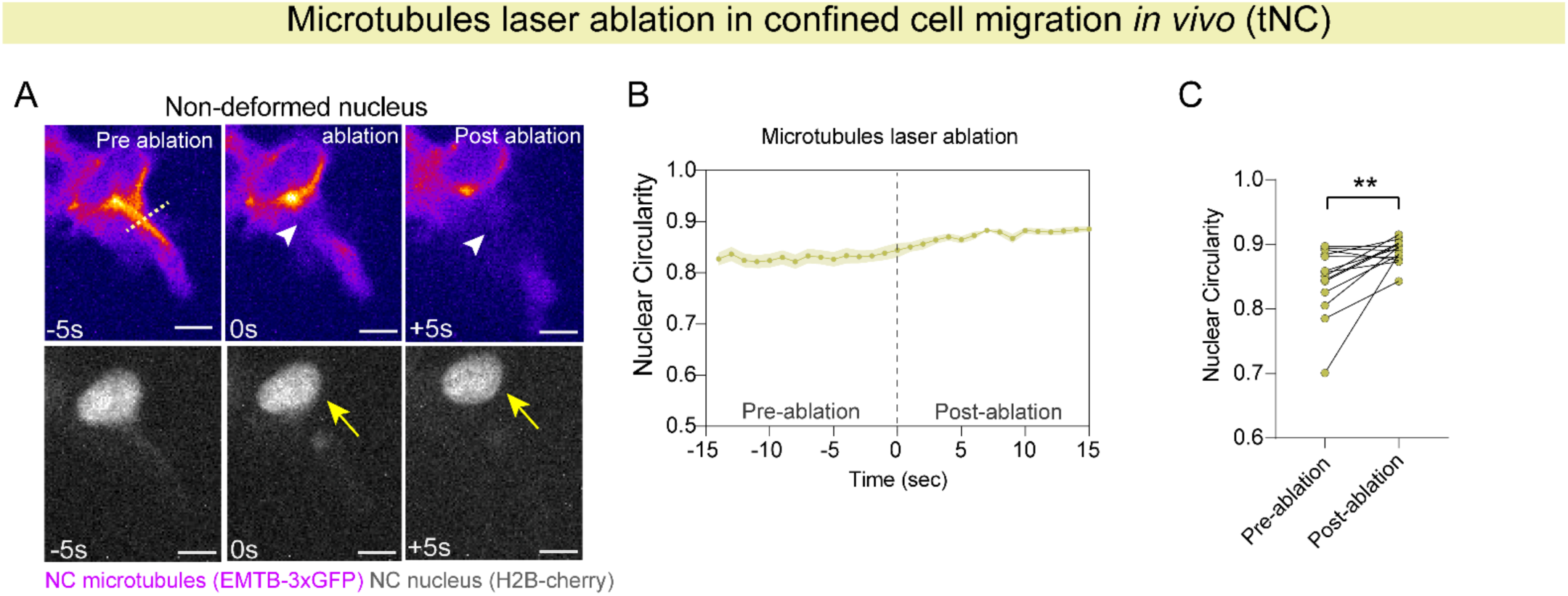
Laser ablation of microtubule bundles also reshapes non-deformed nuclei in confined tNC cells. (A) Representative images of a migrating tNC cell with a non-deformed nucleus during laser ablation (Sox10:Kalt4 / UAS:EMTB-3xGFP). Microtubules are displayed in pseudocolour and the nucleus in grey. White arrowheads indicate the ablation site and yellow arrows indicate nuclear recoil. Scale bar, 5 µm. (B) Mean nuclear circularity over time aligned to the point of microtubule ablation (time = 0). tNC non-deformed nuclei also exhibited recoil and changes in nuclear shape following microtubule severing, consistent with release of mechanical tension. *n* = 14 cells from 14 embryos across 8 experiments. Error bars represent SEM. (C) Quantification of nuclear circularity before and after ablation in tNC cells containing non-deformed nuclei. Two-tailed Wilcoxon matched-pairs signed-rank test; ** *p* = 0.0016. *n* = 14 cells from 14 embryos across 8 experiments.

**Figure. S6.**
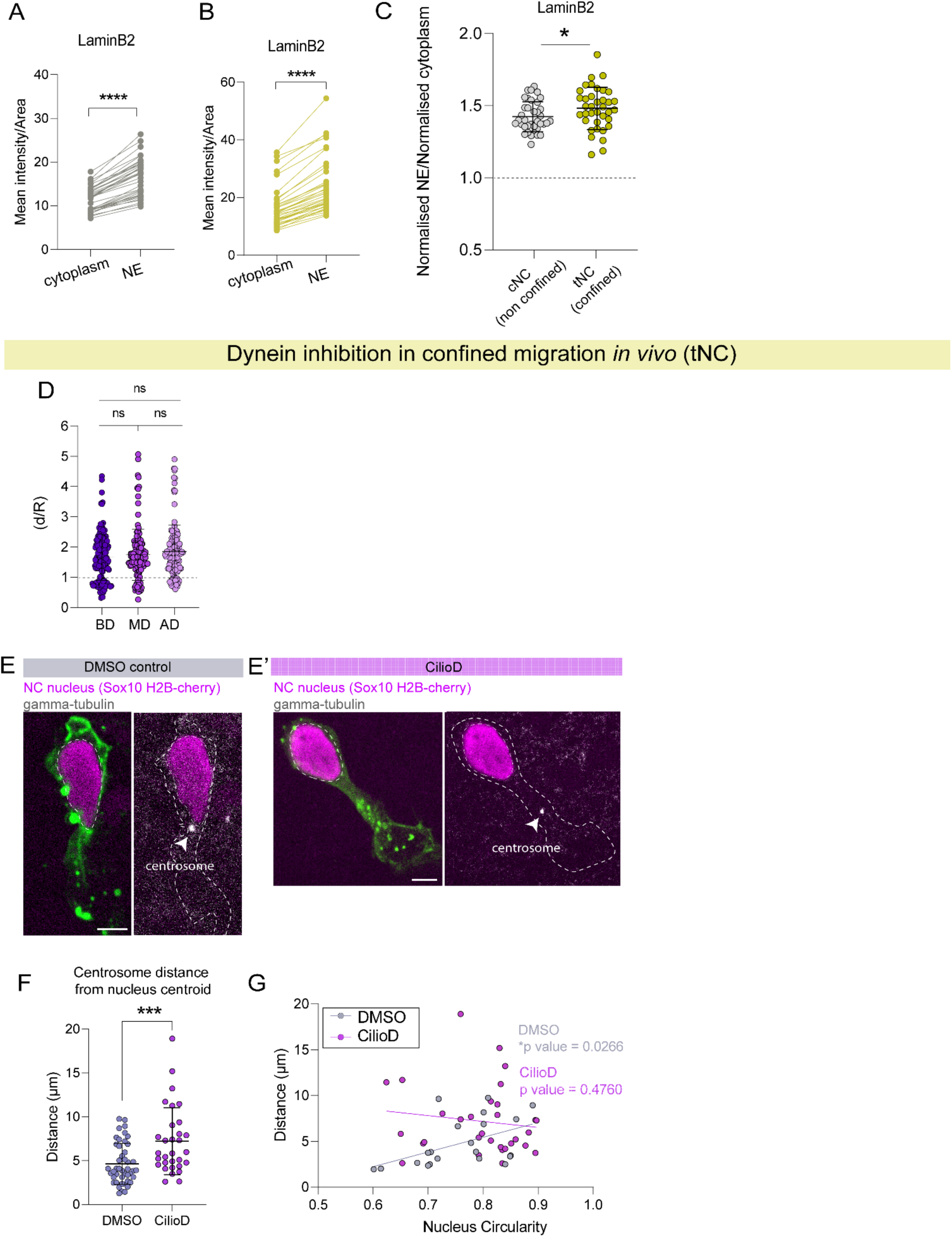
Dynein recruitment to the nuclear envelope correlates with centrosome repositioning during confined migration *in vivo*. (A-B) Quantification of normalized mean intensity per area of Lamin B2 at the nuclear envelope (NE) relative to cytoplasmic regions in non-confined cNC cells (A) and confined tNC cells (B), used as a positive control for nuclear envelope localization. Lamin B2 was significantly enriched at the nuclear envelope in both conditions. Two-tailed Wilcoxon matched-pairs signed-rank test; ****, *p* < 0.0001. cNC: *n* = 37 cells from 6 embryos across 3 experiments. tNC: *n* = 37 cells from 9 embryos across 3 experiments. (C) Ratio of normalized nuclear envelope to cytoplasmic Lamin B2 intensity in non-confined cNC and confined tNC cells as a control for nuclear envelope enrichment. *n* = 74 cells (37 cNC and 37 tNC) from 14 embryos across 3 experiments. Mann–Whitney test; * *p* = 0.0450. (D) Quantification of centrosome-induced nuclear indentation in Ciliobrevin D–treated cells before deformation (BD), at maximum deformation (MD), and after deformation (AD). MD corresponds to the time point of minimum nuclear circularity. No significant differences in nuclear indentation were detected across conditions. *n* = 14 cells comprising 428 total time points (163 BD, 133 MD, and 132 AD) from 8 embryos across 5 experiments. Kruskal–Wallis test with multiple comparisons: BD vs MD (*p* > 0.9999), BD vs AD (*p* = 0.5365), and MD vs AD (*p* = 0.8821). Each dot represents one time point. (E-E’) Representative immunofluorescence images of centrosomes (γ-tubulin, grey) in tNC cells from Sox10:mG embryos (nuclei in magenta, membrane in green) treated with DMSO (E) or Ciliobrevin D (E**′**). In control cells, centrosomes localize closer to the nuclear centroid, whereas dynein inhibition increases centrosome distance from the nucleus. Arrowheads indicate centrosome position. Scale bar, 5 µm. (F) Quantification of centrosome distance from the nuclear centroid in DMSO- and Ciliobrevin D–treated tNC cells. Dynein inhibition significantly increased centrosome distance from the nucleus. DMSO: *n* = 45 cells; CilioD: *n* = 31 cells. Two-tailed Mann–Whitney test; *** *p* = 0.0005. Each dot represents one cell. (G) Linear regression analysis of centrosome distance from the nuclear centroid versus nuclear circularity in fixed tNC cells treated with DMSO (grey) and Ciliobrevin D (purple). In control conditions, increased nuclear circularity (≈1) correlated with greater centrosome distance from the nuclear centroid, whereas increased nuclear deformation (lower circularity) correlated with centrosome proximity to the nucleus. This relationship was significant in DMSO-treated cells (*n* = 45 cells, *p* = 0.0266), but was lost following dynein inhibition (*n* = 31 cells, *p* = 0.4760).

**Figure S7.**
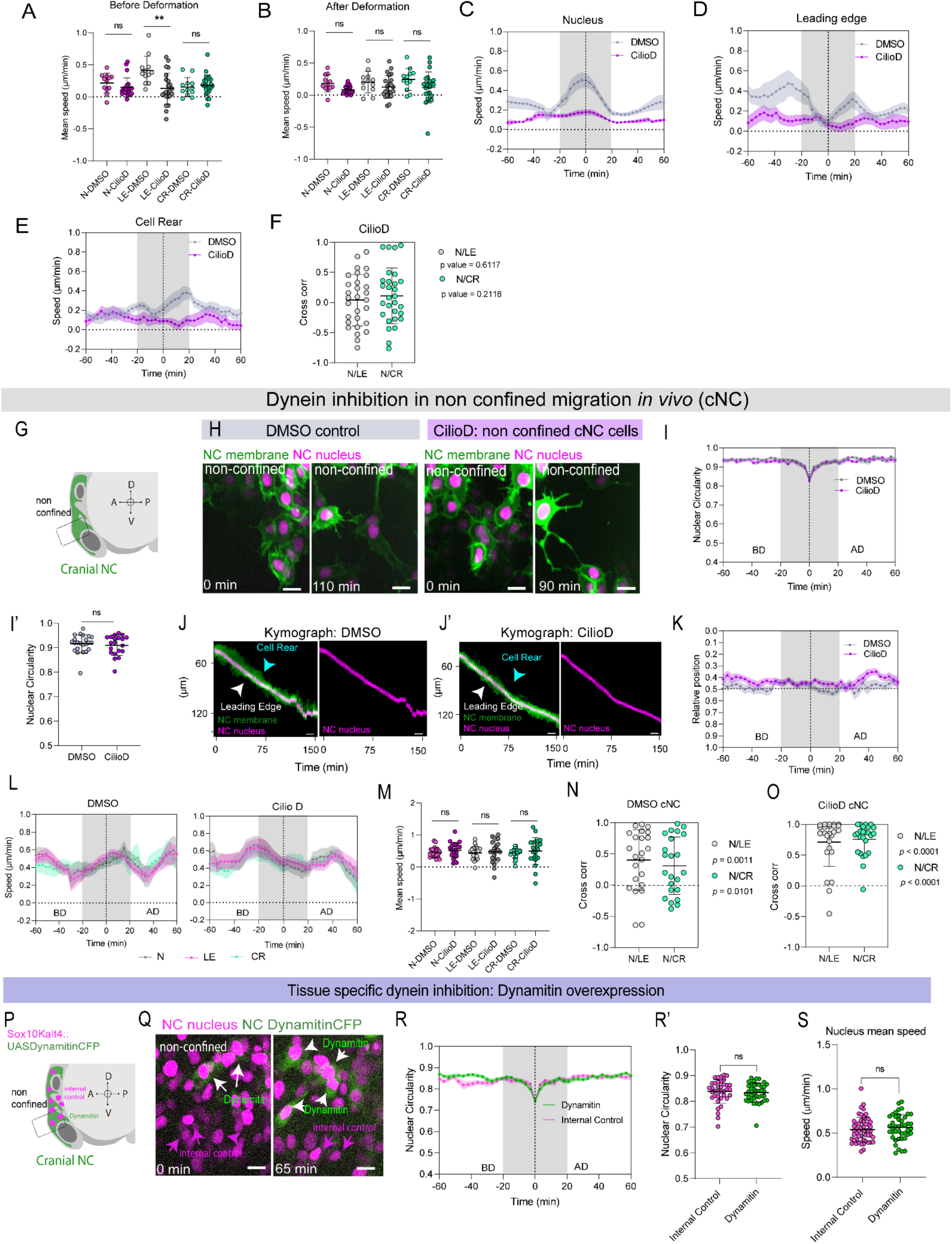
Dynein is dispensable for nuclear translocation and nuclear deformation during non-confined migration *in vivo*. (A) Statistical comparison of mean nuclear, leading-edge (LE), and cell rear (CR) speeds in confined tNC cells before deformation (−60 to −20 min relative to maximum nuclear deformation) between DMSO- and Ciliobrevin D–treated embryos. Kruskal–Wallis multiple comparisons test revealed a significant reduction in leading-edge speed following dynein inhibition (*p* = 0.0024), whereas nuclear and cell rear speeds were not significantly affected (*p* > 0.9999). DMSO: *n* = 12 cells from 4 embryos across 3 experiments; CilioD: *n* = 23 cells from 6 embryos across 3 experiments. Only cells with matched measurements before and after nuclear deformation were included. (B) Statistical comparison of mean nuclear, leading-edge, and cell rear speeds in confined tNC cells after deformation (+20 to +60 min relative to maximum nuclear deformation) between DMSO- and Ciliobrevin D–treated embryos. No significant differences were detected in leading-edge (*p* > 0.9999), nuclear (*p* = 0.5497), or cell rear (*p* = 0.7942) speeds. DMSO: *n* = 12 cells from 4 embryos across 3 experiments; CilioD: *n* = 23 cells from 6 embryos across 3 experiments. Only cells with matched measurements before and after nuclear deformation were included. (C-E) Mean temporal dynamics of nuclear speed (C), leading-edge speed (D), and cell rear speed (E) in DMSO-and Ciliobrevin D–treated confined tNC cells aligned to maximum nuclear deformation (time = 0). BD, before deformation; AD, after deformation. The grey shaded region highlights the period associated with nuclear translocation. Error bars represent SEM. (F) Cross-correlation analysis between nucleus (N) and leading edge (LE), and between nucleus (N) and cell rear (CR), in confined tNC cells treated with Ciliobrevin D. Correlation coefficients were tested against a theoretical value of 0 (no correlation) using a one-sample *t*-test. Values approaching 1 indicate positive correlation, whereas values approaching 0 indicate absence of correlation. Neither N–LE (*p* = 0.6117) nor N–CR (*p* = 0.2118) differed significantly from zero, indicating that dynein inhibition does not restore coupling between nuclear movement and cell edge dynamics. *n* = 27 cells from 6 embryos across 3 experiments. (G) Schematic of cranial neural crest (cNC) migration in a 14-somite-stage zebrafish embryo within a non-confined environment. (H) Representative images of cNC cells migrating in a non-confined environment under DMSO (left) and Ciliobrevin D (right) treatment (membrane in green, nucleus in magenta). Scale bar, 10 µm. (I) Mean nuclear circularity aligned to the point of maximum nuclear deformation (time = 0), spanning 30 min before and after alignment in DMSO- and Ciliobrevin D–treated embryos. DMSO: *n* = 22 cells from 4 embryos across 2 experiments; CilioD: *n* = 21 cells from 4 embryos across 2 experiments. Error bars represent SEM. (I’) Quantification of mean nuclear circularity during the nuclear translocation window (−20 to +20 min relative to maximum nuclear deformation) in cNC non-confined cells from DMSO- and Ciliobrevin D–treated embryos. Each dot represents one cell. Two-tailed Mann–Whitney test; ns, *p* = 0.7632. DMSO: *n* = 22 cells from 4 embryos across 2 experiments; CilioD: *n* = 21 from 4 embryos across 2 experiments. (J-J’) Representative kymographs of membrane and nuclear intensity profiles from DMSO (J) and Ciliobrevin D–treated (J′) cNC cells aligned to the nuclear centroid along the migration axis. Cell rear dynamics are indicated by cyan arrowheads and leading-edge dynamics by white arrowheads. (K) Relative nuclear position within the cell over time in DMSO-treated (grey) and Ciliobrevin D–treated (purple) cNC cells aligned to maximum nuclear deformation (time = 0). Values approaching 0 indicate rear localization, values near 0.5 indicate central positioning, and values approaching 1 indicate localization toward the leading edge. The grey shaded region indicates the period corresponding to nuclear translocation. BD, before deformation; AD, after deformation. DMSO: *n* = 22 cells from 4 embryos across 2 experiments; CilioD: *n* = 21 cells from 4 embryos across 2 experiments. Error bars represent SEM. (L) Mean nuclear speed (N, magenta), leading-edge speed (LE, grey), and cell rear speed (CR, cyan) over time in DMSO (left) and Ciliobrevin D–treated (right) cNC cells aligned to maximum nuclear deformation (time = 0). Data span 60 min before and after alignment. The grey shaded region indicates the analysis window (−20 to +20 min). Dynein inhibition does not alter coordination between nuclear and cell edge dynamics. DMSO: *n* = 22 cells from 4 embryos across 2 experiments; CilioD: *n* = 21 cells from 4 embryos across 2 experiments. Error bars represent SEM. (M) Statistical comparison of mean nuclear, leading-edge, and cell rear speeds during the deformation window (−20 to +20 min relative to maximum nuclear deformation) between DMSO- and Ciliobrevin D–treated cNC cells. Kruskal–Wallis multiple comparisons test revealed no significant differences in leading-edge, nuclear, or cell rear speeds (*p* > 0.9999). DMSO: *n* = 22 cells from 4 embryos across 2 experiments; CilioD: *n* = 21 cells from 4 embryos across 2 experiments. (N-O) Cross-correlation analysis between nucleus (N) and leading edge (LE), and between nucleus (N) and cell rear (CR), in non-confined cNC cells treated with DMSO (N) or Ciliobrevin D (O). Correlation coefficients were tested against a theoretical median of 0 using a one-sample Wilcoxon test. Both N–LE (DMSO: ** *p* = 0.0011; CilioD: **** *p* < 0.0001) and N–CR (DMSO: * *p* = 0.0101; CilioD: **** *p* < 0.0001) differed significantly from zero, indicating maintained coordination between nuclear and cell edge dynamics. (P) Schematic of tissue-specific dynein inhibition in cNC cells using Sox10:Kalt4/UAS:Dynamitin-CFP, induced by tamoxifen. Dynamitin-positive cells display magenta nuclei and green cytoplasm, whereas neighbouring non-expressing cells serve as internal controls.. (Q) Representative images of Dynamitin-expressing cNC cells. White arrowheads indicate Dynamitin-positive cells (green), whereas magenta arrowheads indicate neighbouring control cells. Scale bar, 10 µm. (R) Mean nuclear circularity aligned to maximum nuclear deformation (time = 0) in control and Dynamitin-expressing cells. Internal control: n = 37 cells from 3 embryos across 2 experiments; Dynamitin: *n* = 40 cells from 3 embryos across 2 experiments. Error bars represent SEM. (R’) Quantification of mean nuclear circularity during the nuclear translocation window (−20 to +20 min relative to maximum nuclear deformation) comparing internal controls and Dynamitin-positive cNC cells. Each dot represents one cell. Two-tailed Mann–Whitney test; ns, *p* = 0.3967. (S) Mean nuclear speed in control and Dynamitin-expressing cNC cells. Internal control: *n* = 37 cells, 53 tracks; Dynamitin: *n* = 40 cells, 43 tracks. Two-tailed Mann–Whitney test; ns, *p* = 0.2493.

