## Supplementary Movie Legends for "A dynein-driven nucleokinesis program enables neural crest migration through confined tissues *in vivo*"

**Supplementary Movie 1. Confined tNC cells uncouple nucleokinesis from membrane dynamics in vivo.** Left panel: cNC cells (non-confined). White arrowheads indicate leading-edge extension, while the yellow arrow indicates coordinated movement of the nucleus with membrane dynamics. Right panel: tNC cells (confined). White arrowheads indicate leading-edge extension. Yellow arrows highlight nucleokinesis becoming uncoupled from membrane dynamics. The cyan arrowhead indicates rear membrane retraction. Green: membrane. Magenta: nucleus. Scale bar: 10 μm.

**Supplementary Movie 2. Myosin localization during confined migration of tNC cells.** White arrowheads indicate regions of myosin enrichment (pseudocolour scale), characterized by low myosin accumulation at cell–cell contacts and the leading edge (first pause), and higher accumulation at the rear of the cell (second pause). The yellow arrow indicates the nucleokinesis process during migration. Gray labels the nucleus and cell membrane. Scale bar: 10 μm.

**Supplementary Movie 3. Nucleokinesis occurs independently of ROCK-mediated contractility in confined tNC cells.** Left panel: DMSO-treated embryos. Yellow arrows indicate the nucleokinesis process, and the cyan arrowhead marks membrane retraction at the cell rear. Right panel: embryos treated with the ROCK inhibitor Y-27632. Yellow arrows indicate that nucleokinesis remains largely unaltered despite ROCK inhibition. The white arrowhead highlights the failure of rear membrane retraction in treated cells. Green: membrane. Magenta: nucleus. Scale bar: 10 μm.

**Supplementary Movie 4. StableMark dynamics differ between non-confined cNC cells and confined tNC cells.** Left: non-confined cNC cells. Stable microtubules are labelled with StableMark (green), and nuclei are shown in magenta. White arrowheads indicate stable microtubules in the perinuclear region. Right: confined tNC cells. Stable microtubules are labelled with StableMark (green), and nuclei are shown in magenta. White arrowheads indicate a bundle of stable microtubules at the front of the nucleus during confined migration. Scale bar: 10 μm

**Supplementary Movie 5. EB3-GFP dynamics differ between non-confined cNC cells and confined tNC cells.** Left panel: non-confined cNC cells displaying EB3-GFP growth dynamics (green). Right panel: confined tNC cells displaying EB3-GFP growth dynamics (green). White arrowheads highlight a microtubule bundle forming in front of the nucleus. Magenta labels nucleus and cell membrane. Scale bar: 10 μm.

**Supplementary Movie 6. Centrosome-induced nuclear indentation and deformation during confined migration of tNC cells**. Left panel: non-confined cNC cells displaying EB3-GFP in pseudocolor to highlight centrosome positioning. White arrowheads indicate the variable localization of the centrosome during cNC cell migration, without detectable nuclear indentation or deformation. Right panel: confined tNC cells displaying EB3-GFP in pseudocolor to highlight centrosome positioning. White arrowheads indicate centrosome localization and show how the centrosome initially polarizes to the front of the nucleus. Then moves laterally, where it indents and deforms the nucleus. The nucleus and cell membrane are shown in gray. Scale bar: 10 µm.

**Supplementary Movie 7. Subcellular microtubule laser ablation in non-confined cNC cells does not induce nuclear recoil or deformation.** Left panel: microtubules are displayed in pseudocolor. The white arrow indicates the time point of IR-laser ablation, and white arrowheads mark the recoil of severed microtubules following ablation. Right panel: the nucleus is shown in grayscale. Despite microtubule recoil, no detectable nuclear recoil, deformation, or reshaping is observed following laser ablation. Scale bar: 5 µm.

**Supplementary Movie 8. Subcellular microtubule laser ablation in confined tNC cells reveals nuclear relaxation following microtubule severing (deformed nucleus).** Left panel: microtubules are displayed in pseudocolor. The white arrow indicates the time point of IR-laser ablation, and white arrowheads mark the recoil of severed microtubules following ablation. Right panel: the nucleus is shown in grayscale. White arrowheads indicate nuclear relaxation and reshaping following microtubule severing, consistent with the release of mechanical deformation imposed on the nucleus. Scale bar: 5 µm.

**Supplementary Movie 9. Subcellular microtubule laser ablation in confined tNC cells reveals nuclear recoil following microtubule severing (non-deformed nucleus).** Left panel: microtubules are displayed in pseudocolor. The white arrow indicates the time point of IR-laser ablation, and white arrowheads mark the recoil of severed microtubules following ablation. Right panel: the nucleus is shown in grayscale. White arrowheads indicate nuclear recoil following microtubule severing. Scale bar: 5 µm.

**Supplementary Movie 10. Dynein inhibition uncouples the centrosome from the nucleus during confined migration of tNC cells.** Left panel: DMSO-treated embryos displaying EB3-GFP in pseudocolor to highlight centrosome localization. White arrowheads indicate the centrosome initially polarized at the front of the nucleus before moving laterally and indenting the nucleus. Right panel: embryos treated with Ciliobrevin D (CilioD). White arrowheads indicate that the centrosome initially polarizes anterior to the nucleus but subsequently becomes uncoupled, moving away from the nucleus and failing to induce nuclear indentation, nucleus remains at the back of the cell. The nucleus and cell membrane are shown in gray. Scale bar: 10 µm.

**Supplementary Movie 11**. **Dynein inhibition impairs nucleokinesis in confined tNC cells**. Left: DMSO-treated embryos; yellow arrows indicate nucleokinesis during confined tNC cell migration. Right: CilioD-treated embryos; white arrowheads indicate leading-edge extension, and yellow arrows indicate failed nucleokinesis, with the nucleus remaining at the rear of the cell. Scale bar: 10 μm.

**Supplementary Movie 12. Tissue-specific dynein inhibition impairs nucleokinesis during confined migration of tNC cells.** Cells expressing Dynamitin-CFP are identified by green cytoplasmic fluorescence, whereas Dynamitin-CFP-negative cells are visualized by their magenta-labelled nuclei only. White arrowheads indicate Dynamitin-CFP-positive cells, while magenta arrows mark neighbouring Dynamitin-CFP-negative cells used as internal controls. Dynein inhibition in Dynamitin-CFP-expressing cells impairs nucleokinesis during confined migration. Scale bar: 10 μm.

**Supplementary Movie 13. Dynein inhibition does not impair migration of non-confined cNC cells.** Left panel: DMSO-treated embryos. White arrowheads indicate coordinated movement of the nucleus and cell membrane during non-confined migration. Right panel: embryos treated with Ciliobrevin D (CilioD). White arrowheads indicate that coordinated movement of the nucleus and cell membrane is maintained during non-confined migration. No detectable migration defects are observed following dynein inhibition. Cell membranes are shown in green and nuclei in magenta. Scale bar: 10 μm.

**Supplementary Movie 14. Tissue-specific dynein inhibition does not affect migration of non-confined cNC cells.** Cells expressing Dynamitin-CFP are identified by green cytoplasmic fluorescence, whereas Dynamitin-CFP-negative cells are visualized by their magenta-labelled nuclei only. White arrowheads indicate Dynamitin-CFP-positive cells, while magenta arrows mark neighbouring Dynamitin-CFP-negative cells used as internal controls. No detectable migration defects are observed following dynein inhibition. Scale bar: 10 μm.
